# KIPK and KIPK-LIKE1 suppress overbending during negative hypocotyl gravitropic growth

**DOI:** 10.1101/2024.05.24.595653

**Authors:** Yao Xiao, Melina Zourelidou, Alkistis E. Lanassa Bassukas, Benjamin Weller, Dorina P. Janacek, Lukas Schulz, Sarah Brajkovic, Jan Šimura, Karen Ljung, Bernhard Kuster, Ulrich Z. Hammes, Jia Li, Claus Schwechheimer

**Affiliations:** Plant Systems Biology, School of Life Sciences, Technical University of Munich, Emil-Ramann-Strasse 8, 85354 Freising, Germany; Guangdong Provincial Key Laboratory of Plant Adaptation and Molecular Design, School of Life Sciences, Guangzhou University, Guangzhou, 510006, China; Proteomics and Bioanalytics, School of Life Sciences, Technical University of Munich, Emil-Erlenmeyer-Forum 5, 85354 Freising, Germany; Umeå Plant Science Centre, Department of Forest Genetics and Plant Physiology, Swedish University of Agricultural Sciences, Umeå, Sweden

## Abstract

Plants use environmental cues, such as the direction of gravity or the direction, quantity and quality of light, to orientate organ and plant growth. During germination of angiosperm seeds in the soil, hypocotyl elongation is directed by negative gravitropism responses such that the seedling can reach the light for photosynthesis and autotrophic growth. Hypocotyl elongation in the soil, however, also requires mechanisms to efficiently grow around obstacles such as soil particles. Here, we identify KIPK (KINESIN-LIKE CALMODULIN-BINDING PROTEIN INTERACTING PROTEIN KINASE) and the paralogous KIPKL1 (KIPK-LIKE1) as genetically redundant regulators of hypocotyl bending, in that KIPK and KIPKL1 are required to efficiently align hypocotyl growth with the gravity vector after obstacle avoidance. At the same time, we find that the highly homologous KIPKL2 (KIPK-LIKE2) must be functionally distinct. We further find that KIPK, and likely also KIPKL1, phosphorylate BRXL2 (BREVIS RADIX LIKE2) and ARKs (ARMADILLO REPEAT KINESINs), that mutants of both KIPK phosphorylation substrates share the overbending phenotype with *kipk kipkl1* mutants, and that *KIPK* and *KIPKL1* act synergistically with the ARK-regulatory *NEK6* (*NIMA-RELATED PROTEIN KINASE6*). We propose that KIPK and KIPKL1 regulate ARK kinesins and thereby cortical microtubules for efficient gravitropic hypocotyl bending.

## INTRODUCTION

Plants use environmental cues, such as the direction of gravity or the direction, quantity and quality of light, to orientate organ and plant growth. During germination of angiosperm seeds in the soil, hypocotyl elongation is directed by negative gravitropism. Hypocotyl elongation in the soil, however, also requires mechanisms to efficiently grow around obstacles that impede elongation through the soil (Britz and Galston, 1982; Takahashi and Jaffe, 1990; Takahashi et al., 2000; Gupta et al., 2012). Using the limited resources of the seed, seedling hypocotyls must efficiently combine gravitropic growth and bending responses to reach sunlight for photomorphogenic growth and photosynthesis (Jonsson et al., 2023). Only little is known about how germinating seedlings integrate these responses to direct the hypocotyl through the soil.

The phytohormone auxin regulates many aspects of plant growth and development (Teale et al., 2006). During the directed tropic growth of roots and shoots, differential auxin distribution is commonly observed in the bending tissue, e.g. during negative gravitropic growth, during phototropic bending, as well as during thigmotropic responses, possibly mediated by cell wall loosening and turgor-driven cell elongation (Rakusova et al., 2011; Rakusova et al., 2016; Grones et al., 2018; Lee et al., 2020; Han et al., 2021; Hajny et al., 2022). Indole-3-acetic acid (IAA), the most abundant auxin in land plants, is transported from cell to cell by auxin efflux and influx carriers (Hammes et al., 2022). ‘Canonical’ PIN-FORMED (PIN) auxin transporters are polarly distributed in the plasma membranes of many cells and their polar distribution allows predicting auxin transport through tissues, which may serve to explain auxin-dependent growth, differentiation and tropisms (Teale et al., 2006; Rakusova et al., 2011; Rakusova et al., 2016; Grones et al., 2018; Sauer and Kleine-Vehn, 2019). Tropic root and shoot bending are associated with the differential accumulation of auxin, the presumed result of differential PIN auxin transporter distribution (Friml et al., 2002; Abas et al., 2006). Specifically, *pin2 Arabidopsis thaliana* mutants have agravitropically growing roots (Luschnig et al., 1998), while triple mutants of the paralogous *PIN3*, *PIN4* and *PIN7* (*pin347*) are severely impaired in hypocotyl phototropism and negative gravitropism (Willige et al., 2013). In the hypocotyl, the bending response during negative gravitropism has been explained by the redistribution of PIN3 towards the lower side of the hypocotyl endodermis to allow for auxin accumulation and enhanced cell elongation in the lower side of the hypocotyl (Rakusova et al., 2011). This redistribution is reversed and the bending response is terminated when the optimal angle is reached, presumably as a consequence of the resulting increased cellular auxin accumulation (Rakusova et al., 2016; Grones et al., 2018).

Some members of the ABCB family have also been proposed to act as plasma membrane-resident auxin transporters, but were recently shown to transport brassinosteroid hormones (Noh et al., 2003; Geisler et al., 2005; Geisler et al., 2017; Hammes et al., 2022; Ying et al., 2024). Importantly, single and double mutants of the ABCB transporters *ABCB1/PGP1* and *ABCB19/MDR1/PGP19* show exaggerated hypocotyl bending responses, suggesting that, in the absence of these transporters, the resulting differential auxin or, as more recent findings suggest, brassinosteroid distribution impairs normal tropic growth (Noh et al., 2003).

PIN transporters are activated by AGCVIII family kinases, plant-specific serine/threonine kinases that can be identified by an insertion between protein kinase subdomains VII and VIII (Galvan-Ampudia and Offringa, 2007; Bassukas et al., 2022). In *Arabidopsis thaliana*, the AGCVIII kinases phototropin1 (phot1) and phot2, which form the AGC4 clade of the AGCVIII kinases, are blue light receptor kinases that promote phototropic hypocotyl bending towards the light (Briggs and Christie, 2002). Furthermore, several AGC1 and AGC3 kinases, namely the AGC1 kinases D6 PROTEIN KINASE (D6PK) together with the related D6PK-LIKE1 (D6PKL1) – D6PKL3, as well as the AGC3 kinases PINOID (PID), WAG1 and WAG2 have been implicated in the regulation of photo- and/or gravitropism responses (Sukumar et al., 2009; Rahman et al., 2010; Ding et al., 2011; Willige et al., 2013; Haga et al., 2014). All of the mentioned AGC1 and AGC3 kinases are localized at the plasma membrane where they activate PINs and auxin transport by phosphorylation (Barbosa et al., 2014; Zourelidou et al., 2014). AGC1, but not AGC3 kinases, share the polar distribution with PIN proteins, but their transport to and from the plasma membrane as well as the mechanisms controlling their plasma membrane polarity are distinct from those of PINs (Barbosa et al., 2014; Barbosa et al., 2016; Graf et al., 2024). D6PK and PID plasma membrane interactions are mediated by interactions between polybasic stretches in the kinase insertion domain and anionic phospholipids in the plasma membrane, as well as, in the case of D6PK, S-acylation at conserved repeated CXX(X)P motifs and a phosphoregulation in the insertion domain (Barbosa et al., 2016; Simon et al., 2016; Graf et al., 2024). The biological functions of several AGCVIII kinase family members remain to be established.

Work on the AGC1 kinase PROTEIN KINASE ASSOCIATED WITH BREVIS RADIX (PAX) revealed an important role of BREVIS RADIX (BRX) family proteins in the regulation of auxin transport during protophloem development (Marhava et al., 2018; Marhava et al., 2020; Koh et al., 2021). In this context, BRX emerged as an auxin-labile repressor of PAX, which, at low auxin concentrations, coincides with reduced PAX activation-dependent PIN-mediated auxin transport from the cell and, at high auxin concentrations, leads to increased PAX-dependent auxin export through PINs as a result of BRX inactivation (Marhava et al., 2018; Marhava et al., 2020; Koh et al., 2021). The five-membered family of BRX and BRXL (BRX-LIKE) proteins can be subdivided into two structurally distinct subfamilies and previous work has shown that BRX and BRXL2 are differentially regulated at the cell biological level (Marhava et al., 2018; Marhava et al., 2020; Koh et al., 2021). It remains to be shown whether BRX and the four BRXL proteins have roles also in AGC kinase-dependent processes besides PAX regulation. In this context, it may be relevant that BRXL4 was identified as a regulator of branch angles and gravitropism and interactor of a member of the LAZY protein family (Li et al., 2019; Che et al., 2023).

Kinesin motor proteins interact with microtubules and thereby regulate cellular morphogenesis. *Arabidopsis thaliana* has 61 kinesin-related proteins, which includes ZWICHEL (ZWI), also known as KCBP (KINESIN-LIKE CALMODULIN-BINDING PROTEIN), and Kinesin-13A, regulators of trichome elongation and branching (Oppenheimer et al., 1997; Lu et al., 2005). ARK1 (ARMADILLO REPEAT KINESIN1), ARK2 and ARK3 are three paralogous kinesins that are phosphorylated and regulated by the protein kinase NEK6 (NIMA-RELATED PROTEIN KINASE6), regulators of epidermal cell morphology (Motose et al., 2008; Motose et al., 2011; Zhang et al., 2011; Takatani et al., 2015; Eng et al., 2017; Takatani et al., 2017; Takatani et al., 2020; Lan et al., 2023b; Lan et al., 2023a). NEK6 regulates microtubule responses during hypocotyl elongation in response to tensile stress (Takatani et al., 2020). Research on an ARK ortholog from the moss *Physcomitrium patens* uncovered that ARK is required for the microtubule plus-end-directed anterograde transport of cellular cargos (Yoshida et al., 2023).

Here, we characterize the *Arabidopsis thaliana* AGC1 kinases KIPK (KINESIN-LIKE CALMODULIN-BINDING PROTEIN INTERACTING PROTEIN KINASE, AT3G52890) and its two paralogues KIPKL1/AGC1.9 (KIPK-LIKE1; AT2G36350) and KIPKL2/AGC1.8 (KIPK-LIKE1; AT5G03640) (Galvan-Ampudia and Offringa, 2007). KIPK was originally identified as a yeast two-hybrid interactor of ZWI/KINESIN-LIKE CALMODULIN-BINDING PROTEIN (AT5G65930) but no relevant phenotype has been described with regard to ZWI function in trichome differentiation (Day et al., 2000). An independent study identified KIPK and KIPKL1 as yeast two-hybrid interactors of the putative cell wall-sensing PROLINE-RICH, EXTENSION-LIKE RECEPTOR-LIKE KINASE (PERK10, AT1G26150) and its homologous proteins and mild root growth defects were reported when seedlings were grown on media with high sucrose content (Humphrey et al., 2015). While it is known that the *KIPK* ortholog *DWARF2* (*DW2*) from *Sorghum bicolor* regulates stem internode length (Hilley et al., 2017), as yet, no major biological function has been associated with KIPK/KIPKL proteins in *Arabidopsis thaliana*.

Our study shows that *kipk kipkl1* mutants exhibit exaggerated negative hypocotyl gravitropism when mutants are grown vertically along media and overbending when the seedling hypocotyls grow through soil. We further show that KIPKL2 does not contribute to this phenotype and cannot replace KIPK and KIPKL. Mutants of the reported KIPK interactors *KINESIN-LIKE CALMODULIN-BINDING PROTEIN/ZWICHEL* (*ZWI*) and *PERK* (*PROLINE-RICH EXTENSION-LIKE RECEPTOR-LIKE KINASE*) did not show tropism defects and are unlikely to be involved in negative hypocotyl gravitropism and obstacle avoidance growth examined here. In turn, we find that KIPK/KIPKL proteins interact with and phosphorylate BRX-LIKE proteins and act in concert with ARMADILLO REPEAT KINESIN (ARK) proteins, an interaction, which may allow explaining the exaggerated bending response of the *kipk kipkl1* mutants through the regulation of kinesins and the microtubule network.

## RESULTS

### KIPK, KIPKL1 and KIPKL2 are polarly localized plasma membrane-associated protein kinases

*Arabidopsis thaliana KIPK* and the two closely related *KIPKL1* and *KIPKL2* have unknown biological functions. When compared to other AGC1 kinases including the well-studied D6PK (108 amino acids), these three kinases have extended (537 - 558 amino acids) homologous (27 – 63%) N-termini that are reminiscent of the long N-termini of the AGC4 kinases phot1 and phot2 (576 - 662 amino acids) (Supplementary Figure S1). While these extended N-termini do not contain motifs or domains of known or predictable function, the presence of CXX(X)P repeats in the middle domain and polybasic regions, which, in D6PK, have been shown to function for S-acylation and interactions with plasma membrane phospholipids, respectively, suggests that they may be plasma membrane-associated protein kinases (Barbosa et al., 2016; Bassukas et al., 2022; Graf et al., 2024).

Promoter expression analysis of 2 kb *KIPK*, *KIPKL1* and *KIPKL2* promoter fragments with the GUS (ß-glucuronidase) reporter indicated that all three genes have a broad overlapping expression pattern in dark- and light-grown seedlings and leaves (Supplementary Figure S2). Noteworthy is the speckled staining in root tips, obtained with pKIPKL2::GUS, which is indicative for cell cycle-dependent expression (Supplementary Figure S2C).

To examine the subcellular localization of KIPK and the KIPKL proteins, we expressed YFP- or eGFP-tagged variants from the 2 kb *pKIPK* or *pKIPKL1* promoter fragment from pKIPK::YFP-KIPK, pKIPKL1::eGFP-KIPKL1 or pKIPK::eGFP-KIPKL2. KIPK and the KIPKLs localized polarly at the basal (rootward) plasma membrane in all cells examined, e.g. in all cells of the seeding hypocotyl and root epidermis (Figure 1A, B). The related D6PK is rapidly cycling to and from the plasma membrane, which can be visualized through treatments with Brefeldin A (BFA), an inhibitor of the guanine exchange factor (GEF) GNOM (Figure 1C) (Barbosa et al., 2014). In the case of KIPK, BFA treatment (30 min) had a comparatively minor effect on YFP-KIPK cellular distribution, indicating that KIPK trafficking, unlike D6PK trafficking, may be largely independent from GNOM or slower than D6PK trafficking (Figure 1C).

**Figure 1.**
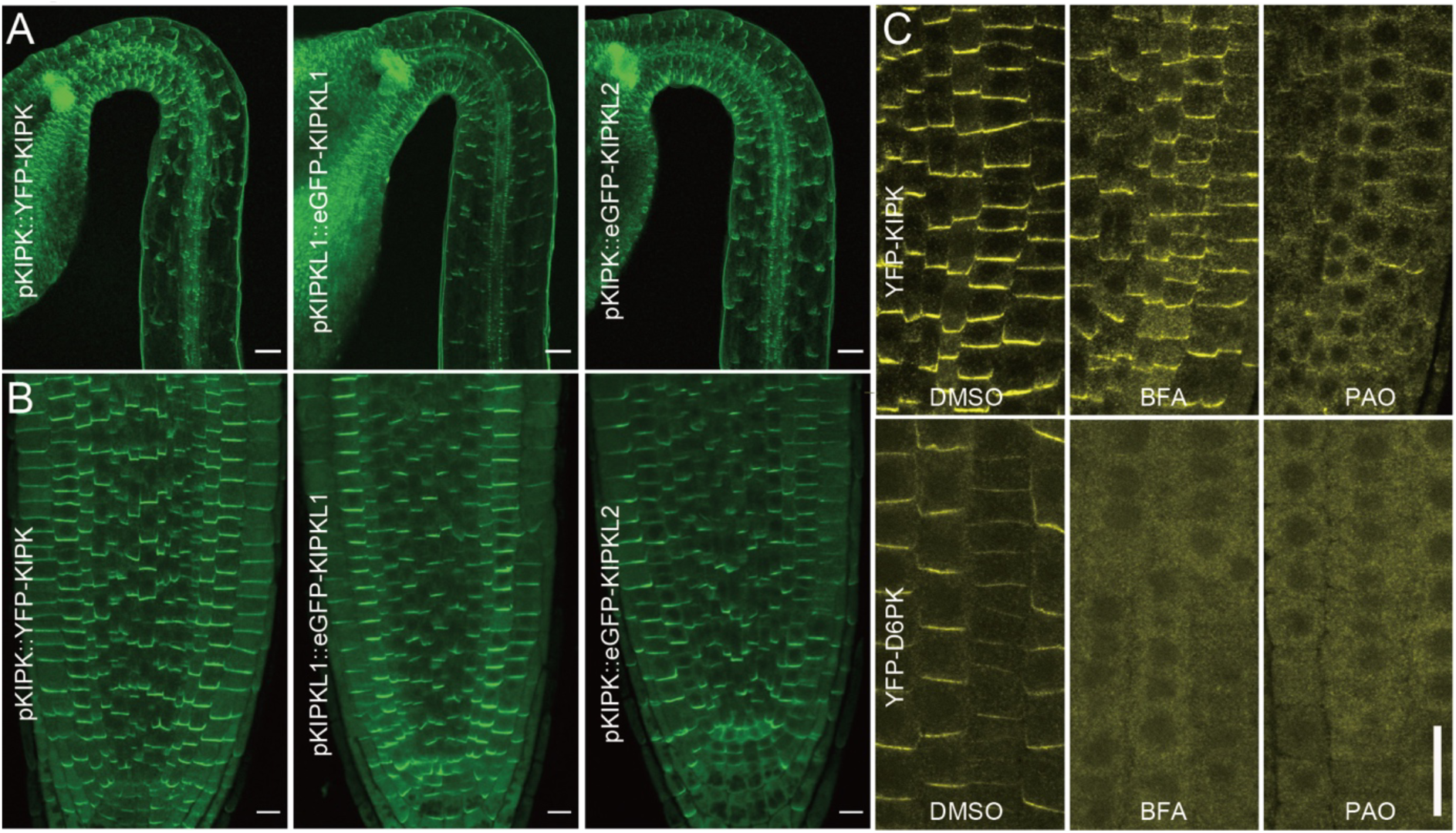
KIPK, KIPKL1 and KIPKL2 are polarly localised plasma membrane-associated protein kinases. **(A)** and (**B)** Representative confocal images of the epidermal cells of hypocotyls with their apical hooks (A) and the root tips (B) from three-days-old dark-grown seedlings expressing the fluorescent protein-tagged KIPK/KIPKLs as specified in the figure panels. Scale bars = 50 µm (A) and 10 µm (B). **(C)** Representative confocal images of root epidermal cells expressing YFP-KIPK and YFP-D6PK following a 30 min mock (DMSO), 25 µM BFA or 30 µM PAO treatment. Note that while YFP-D6PK dissociates fully from the plasma membrane after BFA treatment, YFP-KIPK dissociates only partially under these conditions. Scale bar = 20 µm.

### KIPK, KIPKL1 and KIPKL2 bind anionic phospholipids

Like D6PK and all other AGC1 kinases, KIPK and KIPKLs contain an insertion domain between protein kinase subdomains VII and VIII, which, in the case of D6PK, contains sequence motifs for D6PK plasma membrane association, D6PK recycling and polarity maintenance (Supplementary Figure S1) (Bassukas et al., 2022; Graf et al., 2024). E.g. the D6PK insertion domain contains a polybasic motif required for interactions with negatively charged phospholipids at the plasma membrane and during vesicular trafficking (Barbosa et al., 2016). In experiments with phospholipid-containing PIP strips, recombinant KIPK and KIPKLs bound to mono-, di- and tri-phosphorylated phosphoinositides, as well as to phosphatidic acid, and thereby displayed a similar phosphoinositide preference as D6PK (Supplementary Figure S3A) (Barbosa et al., 2016). Following treatments with a panel of phosphoinositide biosynthesis inhibitors, we identified PAO (phenylarsine oxide), an inhibitor of PtdIns4P (phosphoinositol-4-phosphate) synthesis, to cause dissociation of YFP-KIPK from the plasma membrane (Figure 1C, Supplementary Figure S3B and C)(Simon et al., 2016). Thereby, YFP-KPIK displayed a similar inhibitor sensitivity and behavior as YFP-D6PK with regard to PAO, but dissimilar behavior with regard to U73122, 1-butanol and FIPI, whose effects on YFP-D6PK are, however, less pronounced (Figure 1C; Supplementary Figure S3B, C). We concluded that KIPK required PtdIns4P or its phosphorylated derivatives for interactions with the plasma membrane.

Regions enriched in the basic amino acids arginine (R) and lysine (K), e.g. regions with a basic hydrophobicity (BH) score greater than 0.6, are proposed phospholipid interaction domains of membrane-associated proteins (Bailey and Prehoda, 2015). We identified a highly basic and conserved region in the N-termini of KIPK and KIPKL1 (Supplementary Figure S4A and B) (Bassukas et al., 2022). In this regard, KIPK and KIPKL1 differ from D6PK, which contains a phospholipid binding polybasic region within the insertion domain (Supplementary Figure S4A and C) (Barbosa et al., 2016; Bassukas et al., 2022). Intriguingly, KIPKL2 contained a basic region in the insertion domain, which in turn was absent from KIPK or KIPKL1 (Supplementary Figure S4A and C).

We examined the relevance of the basic region from KIPK for plasma membrane interaction using a KIPK mutant variant where 12 K and R residues were replaced by the uncharged alanine (KR12A) (Supplementary Figure S4B). When expressed as a YFP-fusion, KIPK-YFP (KR12A) localized polarly at the basal plasma membrane and was, in this regard, indistinguishable from wild type YFP-KIPK (Supplementary Figure S4D). We concluded that the highly basic region at the KIPK N-terminus is not required for plasma membrane interactions.

### KIPK and KIPKL1 activate PIN-mediated auxin export in *Xenopus laevis* oocytes

AGC1 (D6PK, PAX) and AGC3 (PID, WAG2) kinases phosphorylate the cytoplasmic loops of ‘long’ PINs and activate PIN-mediated auxin transport from *Xenopus laevis* oocytes (Zourelidou et al., 2014; Marhava et al., 2018). Similarly, recombinant purified MBP- (maltose binding protein-) tagged MBP-KIPK and MBP-KIPKL1 phosphorylated the GST- (glutathione-S-transferase-)tagged PIN3 cytoplasmic loop (GST-PIN3CL) *in vitro*, while GST-PIN3CL phosphorylation by MBP-KIPKL2 was reproducibly comparatively less efficient than phosphorylation by the other two kinases (Figure 2A).

**Figure 2.**
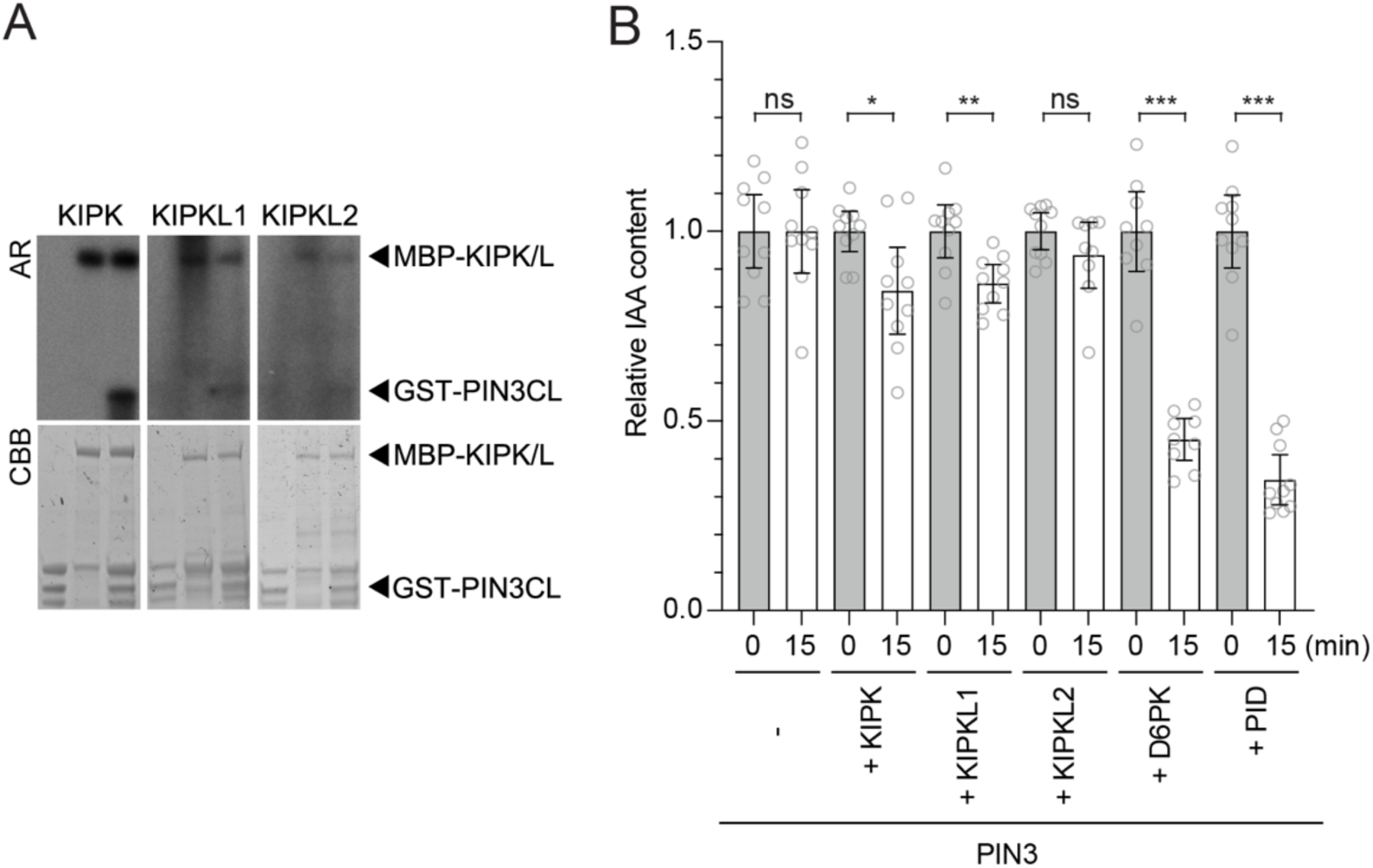
KIPK and KIPKL1 phosphorylate PIN3 and activate PIN3-mediated auxin transport. **(A)** Autoradiographs (AR) and Coomassie Brilliant Blue- (CBB-)stained gels (loading control) from *in vitro* phosphorylation experiments with recombinant purified MBP-KIPK and MBP-KIPKLs (MBP-KIPK/L) and the GST-tagged cytoplasmic loop of PIN3 (GST-PIN3CL). **(B)** Results from auxin transport experiments with PIN3 and protein kinases as specified. Shown is the IAA content after injection (grey bars) set to 1 in comparison to the IAA content after 15 min of efflux (white bars) expressed relative to the starting content. Shown are the individual data points from at least 9 individual oocytes, means and standard errors. Groups were compared by a Student’s t-test: * p < 0.05; ** p < 0.01; *** p < 0.001; ns, not significant.

In *Xenopus laevis* oocytes, PIN3-mediated auxin efflux was activated when *PIN3* was co-expressed with *KIPK* or *KIPKL1*, but non-significantly when co-expressed with *KIPKL2*, suggesting that KIPK and KIPKL1, like D6PK, PAX or PID, are activators of PIN-mediated auxin efflux (Figure 2B)(Zourelidou et al., 2014; Marhava et al., 2018). In comparison with D6PK and PID, controls in our experiment, PIN3 activation by KIPK and KIPKL1 was less efficient, which may be due to differences in kinase activity and activation, protein expression in the oocyte system or, alternatively, a reflection of true biologically relevant differences.

### *kipk01* and *kipk012* mutants display an exaggerated negative gravitropism response

To characterize the biological function of the KIPK and KIPKLs, we isolated homozygous single mutants, *kipk0*, *kipkl1-1*, *kipk1-2*, *kipkl2*, and generated *kipk01* (*kipk0 kipkl1-1*), *kipk01-2* (*kipk0 kipkl1-2*), *kipk02* (*kipk0 kipkl2*), and *kipk12* (*kipk1-1 kipkl2*) double mutants, as well as a *kipk012* (*kipk0 kipk1-1 kipkl2*) triple mutant by genetic crosses (Supplementary Figure S5A). When we examined *kipk012* for the presence of transcript fragments spanning the T-DNA insertion, we did not detect *KIPK* or *KIPKL* transcripts inviting the conclusion that *kipk012* may be a loss-of-function mutant of the three kinases.

The triple mutant did not display apparent defects when grown in the light (Supplementary Figure S5C). Since mutants of other AGCVIII kinases had been implicated in tropic growth responses, we examined gravitropic and phototropic growth in the *kipk and kipkl* mutants. In these experiments, we identified *kipk01* and *kipk012* as mutants with an exaggerated negative gravitropism response in the hypocotyls of dark-grown seedlings when compared to the respective single mutants, the *kipk02* or *kipk12* double mutants or the wild type (Figure 3A - B). Since previous reports had indicated that cotyledon positioning affected the degree of gravitropism response in the hypocotyl, we confirmed and refined mutant phenotyping by resolving gravitropic responses taking into account cotyledon position (Figure 3C) (Khurana et al., 1989). *kipk01* or *kipk012* did not display obvious defects in root gravitropism or phototropic hypocotyl bending, indicating that the bending defect was, at least among the responses tested, specific for negative hypocotyl bending (Figure 3D - E; Supplementary Figure S5C).

**Figure 3.**
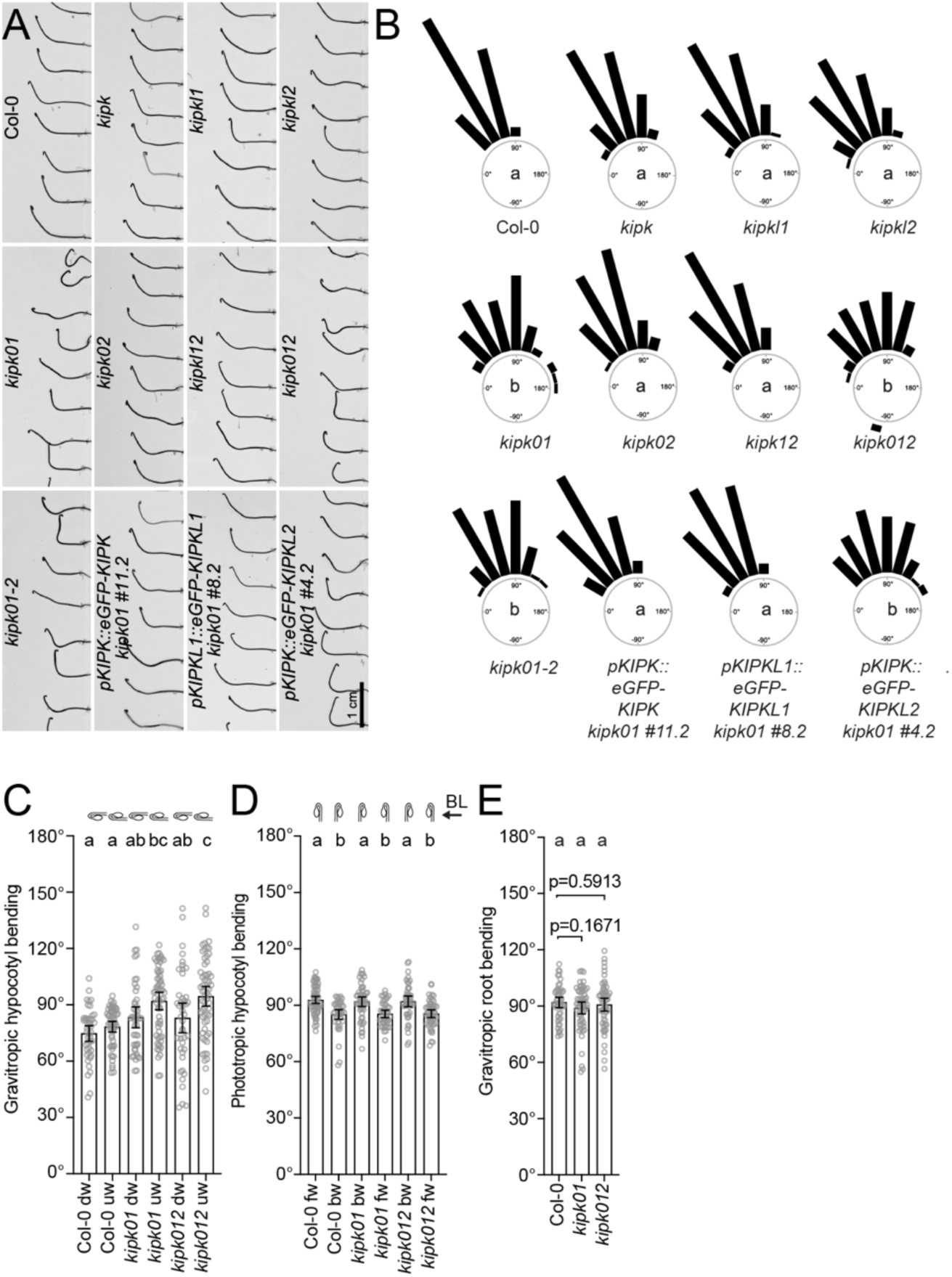
*KIPK* and *KIPKL1* function redundantly in the regulation of negative hypocotyl gravitropism. **(A)** Representative photographs of three-days-old dark-grown seedlings of the specified genotypes 24 hours after reorientation by 90°. **(B)** Rose diagrams displaying gravitropic hypocotyl bending angles of seedlings from the experiment shown in (A). Seedlings were grouped in 15° angle windows. n > 94 seedlings. Results from a One-way ANOVA analysis are displayed in the centres of the diagrams. *kipk01-2* is a second *kipk01* double mutant combination with the *kipkl1-2* allele whose phenotype is indistinguishable from *kipk01-1* carrying the *kipkl1-1* allele used throughout this study. **(C) – (E)** Graphs displaying the average and 95% confidence interval, as well as the individual data points from a negative hypocotyl gravitropism experiment (C) as shown in (A) and (B), a hypocotyl phototropism experiment (D) and a root gravitropism experiment (E). Since it is known that cotyledon positioning influences the degree of hypocotyl bending (Khurana et al., 1989), seedling responses were evaluated independently for seedlings with downward- (dw-) or upward- (uw-) positioned cotyledons (C) or with forward- (fw-) or backward- (bw-) oriented cotyledons with regard to the gravity vector (C) or the orientation of the blue light (BL) used for seedling illumination (D). n > 41 seedlings. Results from a One-way ANOVA analysis are displayed on the top of each bar.

The hypocotyl gravitropism defect of the *kipk01* double mutant was suppressed after expression of *KIPK* or *KIPKL1* from a 2 kb *pKIPK* promoter fragment, but not after expression of *KIPKL2* from *pKIPK* (Figure 3A, B). This suggested that KIPKL2 was biochemically different from the other two family members, a finding also in line with the observation that the loss of *KIPKL2* in the *kipk012* triple mutant did not enhance the hypocotyl gravitropism defect of the *kipk01* double mutant (Figure 3A). Further, expression of the KIPK(KR12A) variant could still suppress the *kipk012* phenotype, suggesting that the mutated motif is not important for KIPK function (Supplementary Figure S4E). In summary, we concluded that *KIPK* and *KIPKL1* are required for the regulation of proper negative gravitropism in *Arabidopsis thaliana* and that *KIPKL2* does not contribute to this process. The observation that *KIPKL2* is unable to complement the *kipk01* phenotype aligned with the differential activity of KIPKL2 in the *in vitro* phosphorylation and auxin transport experiments (Figure 2).

### *kipk01* overbending responses require PIN-mediated auxin transport

The phototropic and gravitropic bending of seedling hypocotyls requires the functionally redundant auxin efflux carriers PIN3, PIN4, and PIN7, as well as their phospho-regulation by D6PK, D6PKL1 and D6PKL2 (Willige et al., 2013). We found that similar to *d6pk d6pkl1 d6pkl2* (*d6pk012*) or *pin3 pin4 pin7* (*pin347*) triple mutants, *kipk01* mutants displayed a strong reduction in basipetal auxin transport in hypocotyls of dark-grown seedlings, suggesting that KIPK and KIPKL1 participate in basipetal auxin transport in Arabidopsis hypocotyls, possibly as a result of the role of KIPK and KIPKL1 in PIN activation (Figure 3, Figure 4A). Following treatments with NPA, an inhibitor of PIN-mediated auxin transport, we found that auxin transport inhibition suppressed the gravitropic responses of the *kipk01* mutants with a similar sensitivity as observed in the wild type or in *pin3* single mutants, which are only partially impaired in negative gravitropism (Figure 4B) (Abas et al., 2021). Introduction of the *pin347* triple mutant into *kipk01* suppressed the gravitropic bending in *kipk01*, a consequence of PIN-mediated auxin transport being required for the initial bending response (Figure 4C). We concluded that PIN-mediated auxin transport is compromized in *kipk01* mutant hypocotyls and that *kipk01* hypocotyl overbending does not supersede the requirement of PIN-mediated auxin transport for bending.

**Figure 4.**
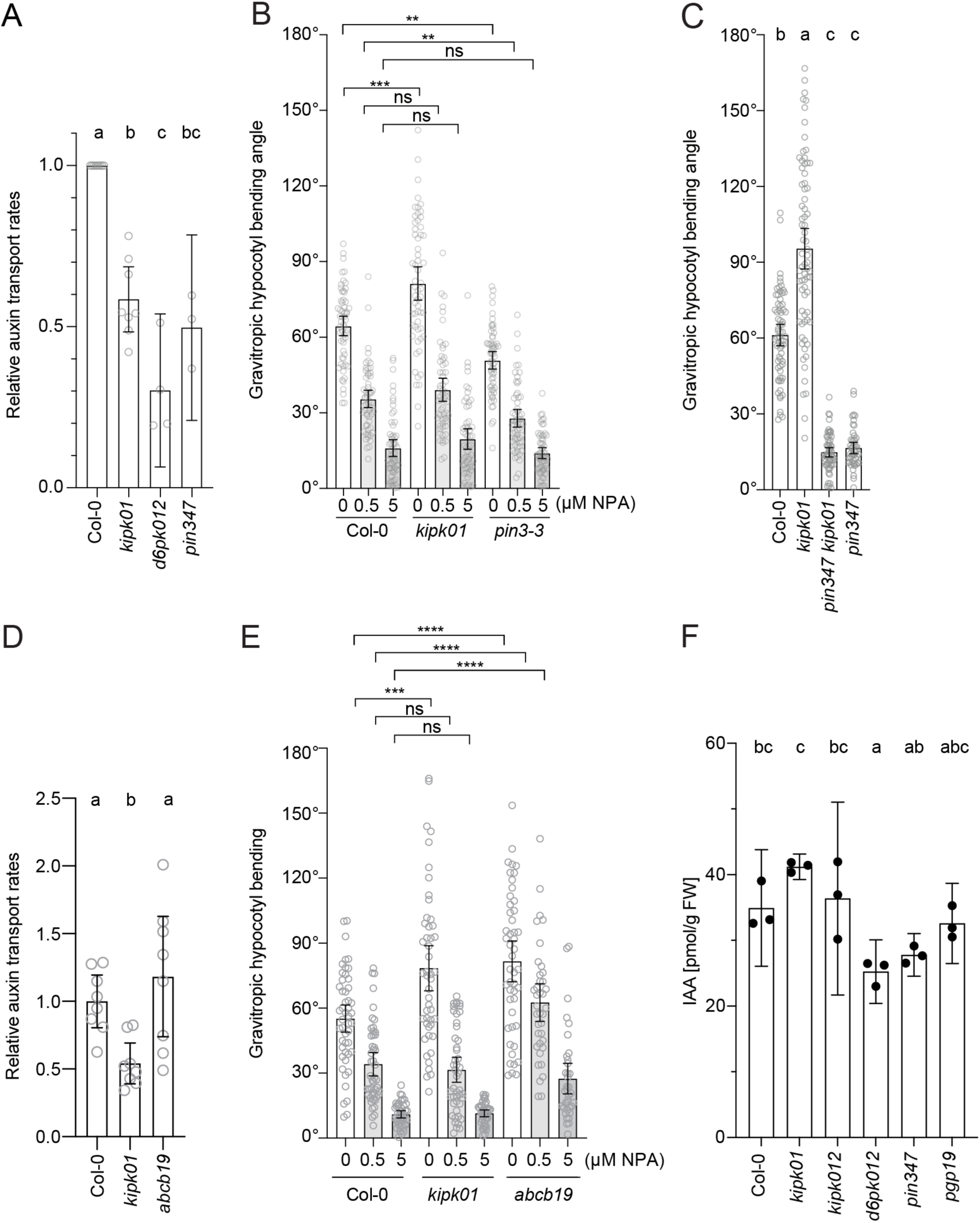
PIN-mediated auxin transport is a prerequisite for *kipk01* hypergravitropic responses. **(A)** and **(D)** Graphs displaying the average and 95% confidence interval, as well as the individual results, of relative auxin transport rates in wild type (Col-0), *kipk01*, *d6pk012*, and *pin347* mutants; n ≥ 3 independent experiments (A) or *kipk01* and *abcb19* mutants n = 8 × 5 seedlings (D). **(B)**, **(C)** and **(E).** Graphs displaying the average and 95% confidence interval, as well as the individual results, of gravitropic hypocotyl bending experiments with three-days-old dark-grown wild type (Col-0), *kipk01* and *pin3-3* seedlings after treatments with NPA (B), with three-days-old dark-grown wild type (Col-0), *kipk01*, *pin347*, *pin347 kipk01* seedlings (C) or with three-days-old dark-grown wild type (Col-0), *kipk01*, *abcb19* seedlings after treatments with NPA (E). n > 37 seedlings. Results from a One-way ANOVA analysis or Student’s t-tests are displayed. Groups were compared by a Student’s t-test: * p < 0.05; ** p < 0.01; *** p < 0.001; ns, not significant. **(F)** Graph displaying the average and standard deviation of IAA quantifications in hypocotyls and cotyledons of pooled four-days-old dark-grown seedlings of the specified genotypes (n = 3 pools). Results from a One-way ANOVA analysis are displayed.

Hypocotyl overbending was previously reported for the ABCB transporter mutant *abcb19* whose phenotype is reminiscent of the phenotype of *kipk01* mutants (Noh et al., 2003). The *abcb19* mutant had, however, a similar auxin transport rate as the wild type and was comparatively less sensitive to NPA than the wild type or the *kipk01* mutant (Figure 4D and E). The *abcb19* mutant has thus defects in auxin transport that are distinct from those of the *kipk01* or *pin347* mutants. While the auxin (IAA) concentrations in hypocotyls and cotyledons were reduced in *d6pk012* and *pin347* mutants, they were comparable between the wild type, *abcb19*, as well as *kipk01* and *kipk012*, indicating that alterations of IAA content may not allow explaining *kipk01* and *kipk012* mutant phenotypes (Figure 4F).

### Auxin accumulation patterns but not KIPK or PIN3 polar distribution change during gravitropic bending

The differential distribution of PIN3 during gravitropic bending between the lateral plasma membranes in hypocotyl endodermis cells had previously been reported (Rakusova et al., 2011; Rakusova et al., 2016). Accordingly, the differential cellular auxin accumulation resulting from differential PIN3 polarity would lead to a redistribution of PIN3 to terminate the bending response (Rakusova et al., 2011; Rakusova et al., 2016; Grones et al., 2018). We reasoned that defects in the above-summarized differential PIN3 distribution may be a causal for the observed bending defects in *kipk01* mutants.

Our assumption that the critical response may be sensed by the endodermis found support in our observation that expression of eGFP-KIPK or mCitrine-KIPK from the endodermis-specific *SCR* (*SCARECROW*) promoter was sufficient to rescue the bending and the auxin transport defects of *kipk01* (Figure 5A - C). Whereas we did not observe any changes in the cellular distribution of eGFP-KIPK or mCitrine-KIPK during gravitropic bending, we noted pronounced accumulations of auxin in *kipk01* mutants, as detected with the auxin response reporter DR5V2::GUS (Figure 5D, Figure 6). In the wild type and in the *kipk01* mutants, these auxin accumulations coincided with the bending regions within the tissues (Figure 6). However, whereas the wild type readily stained for the GUS reporter (4 hrs), staining times had to be extended for *kipk01* (24 hrs) due to the weak DR5V2::GUS staining in the *kipk01* hypocotyls (Figure 6). The latter was accompanied by a strong DR5V2::GUS staining in the *kipk01* cotyledons, which again may be a reflection of the reduced auxin transport observed in the mutants (Figures 4D and 6). We concluded that the overbending of *kipk01* mutants, despite of the mutants having reduced auxin transport, correlated with differential auxin accumulation in the bending regions, suggesting that tissue level auxin transport may be mis-regulated, but not defective in the mutants. The strong auxin accumulation in the cotyledons, as suggested by their strong DR5V2::GUS staining, may be a result of auxin being produced in the cotyledons not being efficiently transport through the hypocotyl, as suggested by our auxin transport studies (Figure 4A, Figure 6). The difference in the responsiveness observed with DR5V2::GS between the wild type and *kipk01* was also observed with other auxin reporter constructs (DR5::GUS, DR5::GFP) when introduced into *kipk01* and may, particularly in view of the comparable auxin contents measured in the mutant and the wild type, be indicative for effects of the *kipk01* mutations on the cellular auxin response machinery (Figure 4F, Figure 6).

**Figure 5.**
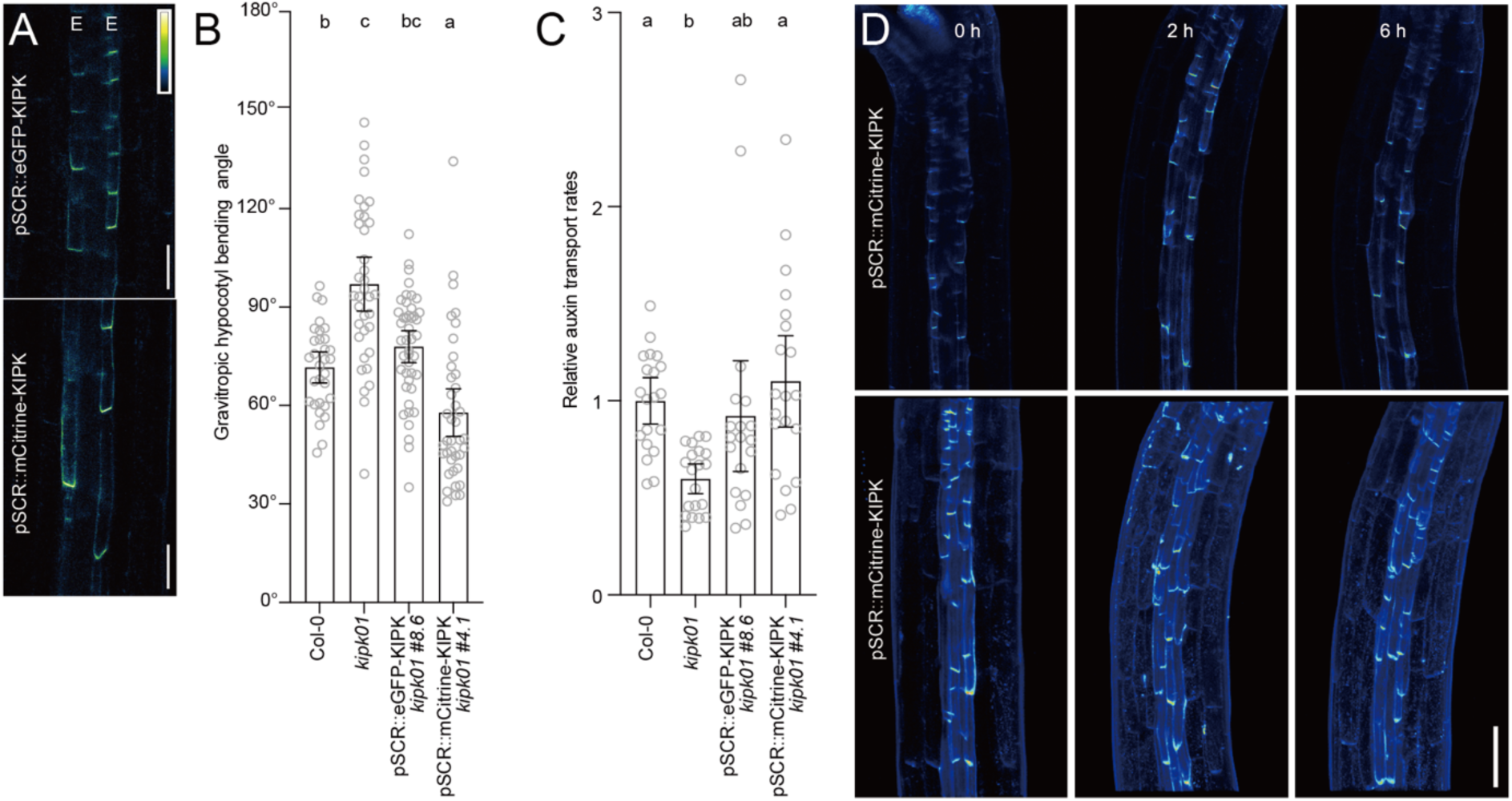
Endodermis-specific expression of *KIPK* suppresses the *kipk01* hypergravitropic responses. **(A)** Representative confocal microscopy images of hypocotyls of three-days-old dark-grown seedlings expressing pSCR::eGFP-KIPK or pSCR::mCitrine-KIPK. E, endodermal cell file. Scale bars = 50 µm. **(B)** Graph displaying the average and 95% confidence interval, as well as the individual results, of gravitropic hypocotyl bending experiments with three-days-old dark-grown wild type (Col-0), *kipk01* and *kipk01* pSCR::eGFP-KIPK or *kipk01* pSCR::mCitrine-KIPK seedlings. n > 31 seedlings. Results from a One-way ANOVA analysis are displayed. **(C)** Graph displaying the average and 95% confidence interval, as well as the individual results, of relative auxin transport rates in wild type (Col-0), *kipk01* and *kipk01* pSCR::eGFP-KIPK or *kipk01* pSCR::mCitrine-KIPK seedlings. n = 20 × 5 seedlings. Results from a One-way ANOVA analysis are displayed. **(D)** Representative confocal images showing the distribution of mCitrine-KIPK before and after gravity stimulation for the times specified in the image in the central axial plane of the hypocotyl (upper images) and in the hypocotyl half cylinder (lower images; z stack). Scale bar = 100 µm.

**Figure 6.**
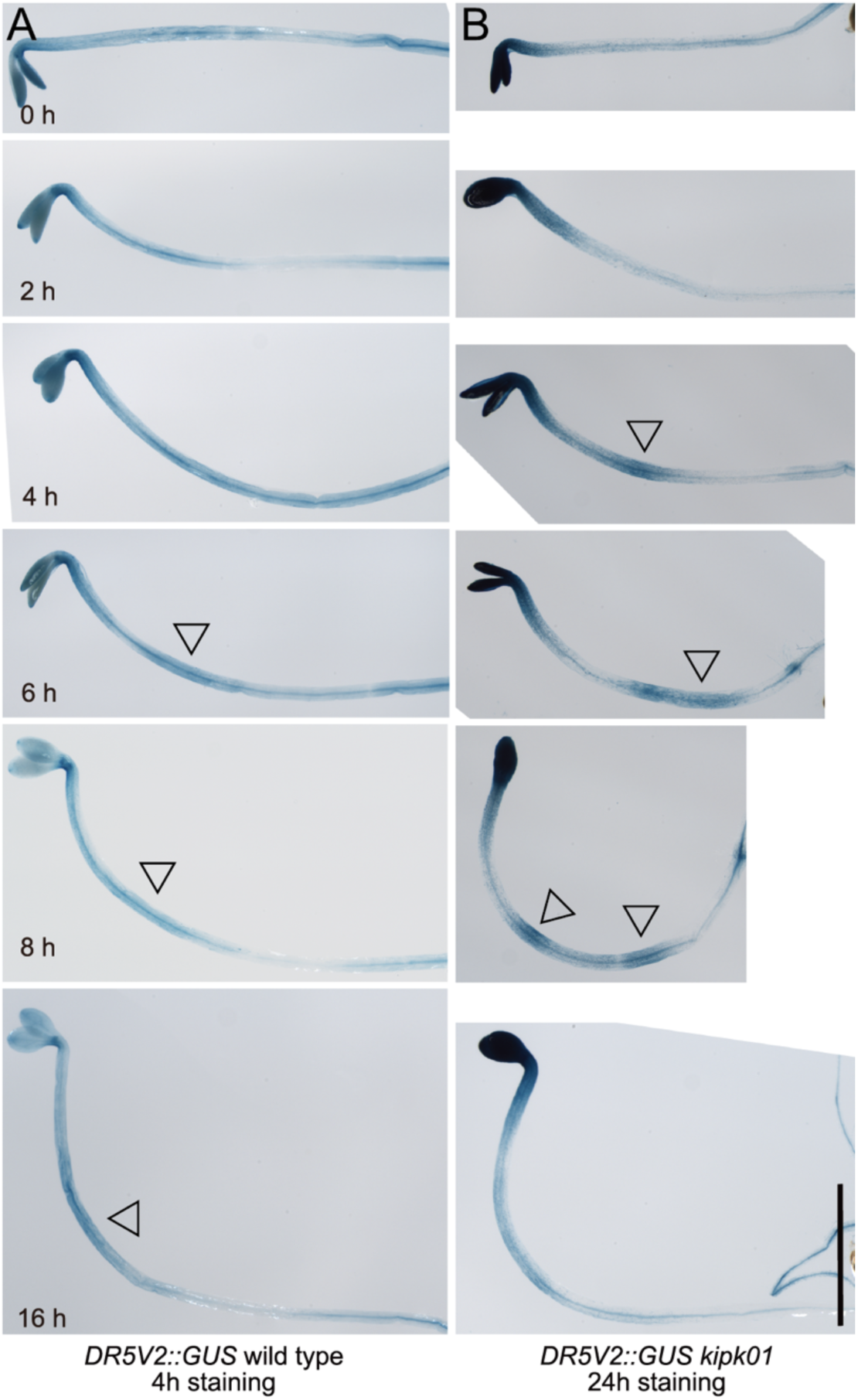
Hypocotyl overbending in *kipk01* coincides with the strong tissue level accumulation of auxin as monitored with DR5V2::GUS. **(A)** and **(B)** Representative photographs of dark-grown seedlings that had been exposed, for the times specified in the image, to a 90° change in the gravitropic vector when they were three days old. Arrowheads mark the sites of auxin accumulation. Note the strong accumulation of auxin in the *kipk01* cotyledons. Scale bar = 5 µm.

To be able to examine PIN3 distribution selectively in the gravity-sensing hypocotyl endodermis cells, we established lines for the expression of PIN3-GFP from the *SCR* promoter. When we measured PIN3-GFP lateral distribution by establishing the ratio between the outer (epidermis-facing) and inner (stele-facing) lateral membrane in lower, as well as the upper endodermal cells, we did not detect a differential distribution of PIN3-GFP during the gravitropic response, a finding that is in conflict with previously reported findings (Figure S6). From these data, we concluded that, at least in our experimental conditions, a lateral redistribution of PIN3 cannot explain gravitropic hypocotyl bending in the wild type or the overbending defect of the mutants. We further employed an anti-GFP antibody to monitor the distribution of PIN3-GFP between the wild type and the *kipk01* mutant. However, these experiments did not reveal any changes in the abundance of PIN3-GFP or in the abundance of a phosphorylated PIN3-GFP form that can be separated on the respective immunoblots between the wild type and the *kipk01* mutants (Figure S6C). We concluded that dynamic changes in the distribution of PIN3-GFP, its abundance or phosphorylation cannot be correlated with the exaggerated hypocotyl bending phenotype of *kipk01* mutants.

### BRXL2 interacts with KIPK in the regulation of hypocotyl gravitropism

To identify KIPK interactors, we performed immunoprecipitation of fluorescent protein-tagged KIPK from protein extracts prepared from dark-grown seedlings. Following mass spectrometry, we identified about 1700 putatively interacting proteins (Figure 7A; Supplementary Data Set S1). 653 of these were specifically enriched in the KIPK purification, and not present in an immunoprecipitation of unfused eGFP-GUS (Figure 7A; Supplementary Data Set S1). Using co-expression analysis at ATTED-II (https://atted.jp/), we found 93 *KIPK* and *KIPKL1* co-expressed genes, nine of which were also retrieved in the KIPK immunoprecipitations but not in the control eGFP-GUS purification (Figure 7A; Supplementary Data Set S1).

**Figure 7.**
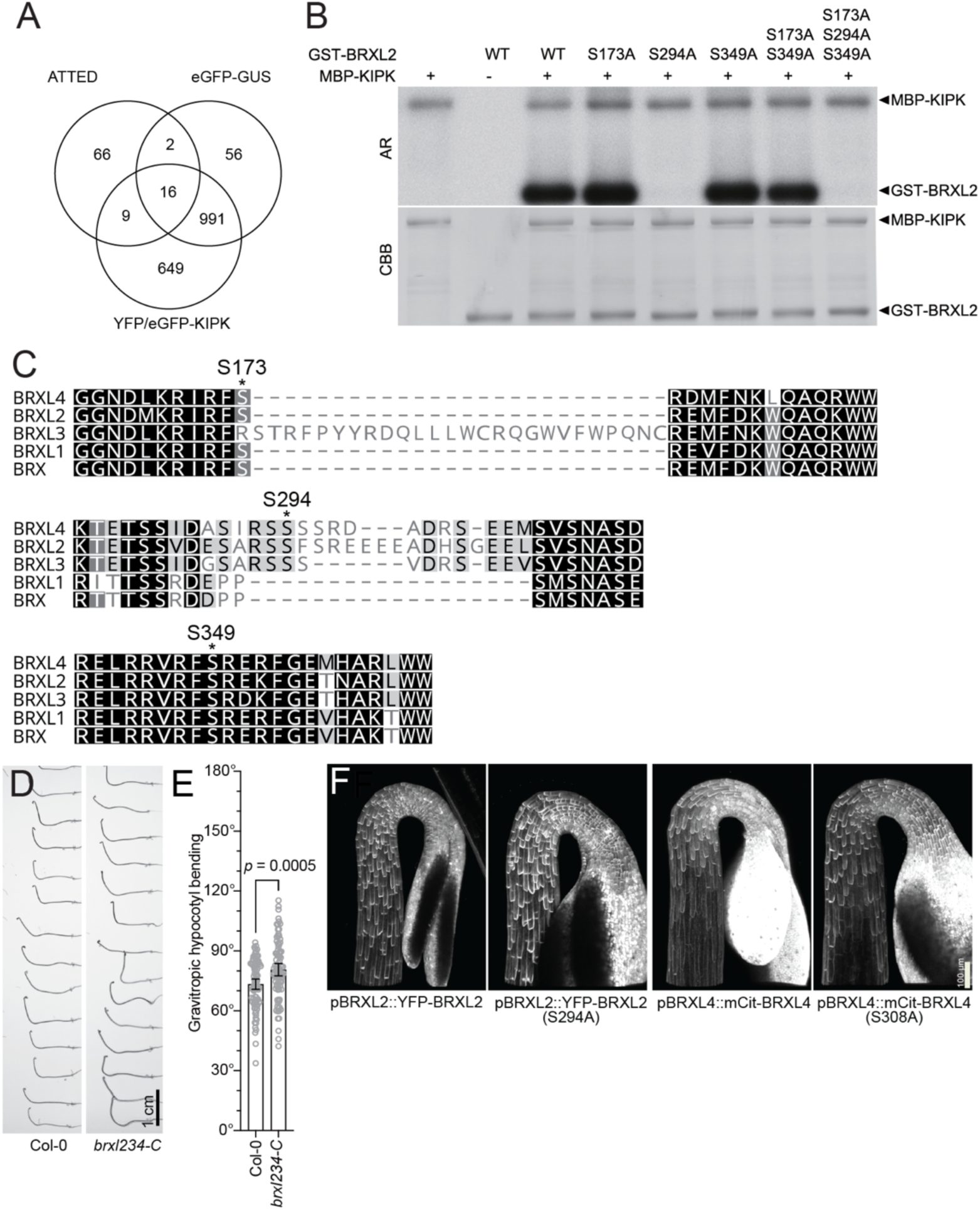
KIPK interacts with BRXL proteins. **(A)** Venn diagram displaying the results from immunoprecipitations of eGFP-GUS-KIPK and the sum of two independent immunoprecipitations of YFP- and GFP-KIPK and the genes identified in an ATTED coexpression analysis (Supplemental Table S1). **(B)** Autoradiograph (AR) and Coomassie Brilliant Blue (CBB) stained gel of an SDS-PAGE with the result of *in vitro* phosphorylation reactions with MBP-KIPK and wild type GST-BRXL2 (WT) and mutant variants with alanine (A) replacement mutations of selected serines (S). **(C)** Pretty box sequence alignment of protein fragments of the five Arabidopsis BRX/BRXL proteins with serines mutated in (B) and (E) highlighted. **(D)** Representative photographs of three-days-old dark-grown seedlings of the specified genotypes 24 hours after reorientation by 90°. **(E)** Graph displaying the average and 95% confidence interval, as well as the individual data points and the result from a Student’s t-test from a negative hypocotyl gravitropism experiment as shown in (D). **(F)** Representative confocal microscopy images of three-days-old dark-grown seedlings expressing the transgenes as specified. Scale bar = 100 µm.

BRXL2 was one of the nine co-expressed immunoprecipitated proteins (Figure 7A; Supplementary Data Set S1). BRX family proteins had previously been shown to interact and to be regulated by the AGC1 kinase PAX (Marhava et al., 2018; Koh et al., 2021). Consistently, we found that GST-tagged BRXL2 can be phosphorylated by recombinant MBP-KIPK (Figure 7B). Based on the known phosphorylation preference of AGC1 kinases for RXS sequence motifs, we examined BRXL2 S173, S294, S349 as three putative phosphorylation sites. We identified BRXL2 S294 as the major KIPK phosphorylation site since a GST-BRXL2 S294A alanine-replacement mutation could not be phosphorylated *in vitro* (Figure 7B and C). BRXL2 S294 is only conserved between the three paralogous BRXL2, BRXL3 and BRXL4, which form a distinct subfamily among the five BRX/BRXL proteins (Figure 7C). We argued that the regulation of KIPK on these proteins may be restricted to this subfamily or possibly only to BRXL2. When we performed a phenotype analysis with a published *brxl2 brxl3 brxl4 (brxl234-T)* mutant, we realized that the T-DNA insertion allele *brxl2* (SALK_032250) was a knock-down rather than a loss-of-function mutant (Supplementary Figure S7) (Briggs et al., 2006). We therefore generated a new *brxl2 brxl3 brxl4* triple mutant using CRISPR/Cas9, *brxl234*-C (Supplementary Figure S7). When we analysed *brxl234*-C for defects in gravitropism response, we found that *brxl234*-C shared the gravitropism defect with the *kipk01* mutants, indicating that the BRXL proteins participate in the KIPK and KIPKL1-dependeng bending control (Figure 7D and E).

YFP-tagged BRXL2 or BRXL4, when expressed from *BRXL2* or *BRXL4* promoter fragments, respectively, shared the strong polar distribution at the basal plasma membrane of all cells of the dark-grown seedling hypocotyl with KIPK, suggesting that the two proteins act in close proximity to each other (Figure 7F). Replacing S294 in pBRXL2:YFP-BRXL2 or S308 in pBRXL4::mCitrine-BRXL4 by alanine did not affect the polar distribution of the proteins at the plasma membranes (Figure 7F).

To examine the potential interplay between BRXL2 and KIPKs in the regulation of auxin transport, we tested the effect of BRXL2 on KIPK-activated PIN-mediated auxin transport in *Xenopus laevis* oocytes. Co-expression of BRX had previously been shown to attenuate the auxin transport activation of D6PK and PAX on PIN-activated auxin transport (Marhava et al., 2018). However, BRXL2 had no noteworthy effect on PIN activation by neither KIPK, nor D6PK or PID, which is consistent with previous findings that BRXL2 is functionally distinct from BRX (Supplementary Figure S8)(Koh et al., 2021).

### *KIPK* and *KIPKL* genetically interact with the ARK kinesin regulatory serine/threonine kinase *NEK6*

*nek6* mutants, defective in the ARK kinesin-regulatory NEK6 (NIMA-RELATED MICROTUBULE-ASSOCIATED KINASE6) serine/threonine kinase, display a hypocotyl overbending defect similar to the one reported here for *kipk01* mutants (Sakai et al., 2008; Takatani et al., 2020). *nek6* mutants display an enhanced sensitivity to microtubule-stabilizing drug taxol and we detected increased taxol sensitivity also in *kipk01* mutants (Figure S9A). We tested the genetic interaction between *KIPK, KIPKL1* and *NEK6* by crossing the *kipk01* double mutant with the *nek6-1* single mutant. Among 100 F3 progeny seedlings from a *kipk* (−/−) *kipkl1* (−/−) *nek6-1*(+/−) mother plant, we identified only *kipk* (−/−) *kipkl1* (−/−) *nek6-1*(+/−) and *kipk* (−/−) *kipkl1* (−/−) *NEK6* plants but no *kipk01 nek6-1* triple mutants. *kipk01 nek6-1* +/− plants displayed a slightly more compact plant rosette phenotype than the *kipk01* or *nek6-1* mutants and produced shorter siliques (Figure S9B). Although we could obtain progeny seed from the *kipk01 nek6-1* +/− segregating line, the siliques contained many aborted seeds, indicating that embryogenesis or seed formation may have been interrupted following fertilization (Supplementary Figure S9C). We subsequently tested negative hypocotyl gravitropism responses in the mutants and confirmed the *nek6* overbending phenotype also in our experimental conditions (Figure 8A and B). The strong defect in hypocotyl bending in progreny seedlings from *kipk* (−/−) *kipkl1* (−/−) *nek6-1*(+/−) was accompanied by strongly reduced hypocotyl elongation (Figure 8A – C).

**Figure 8.**
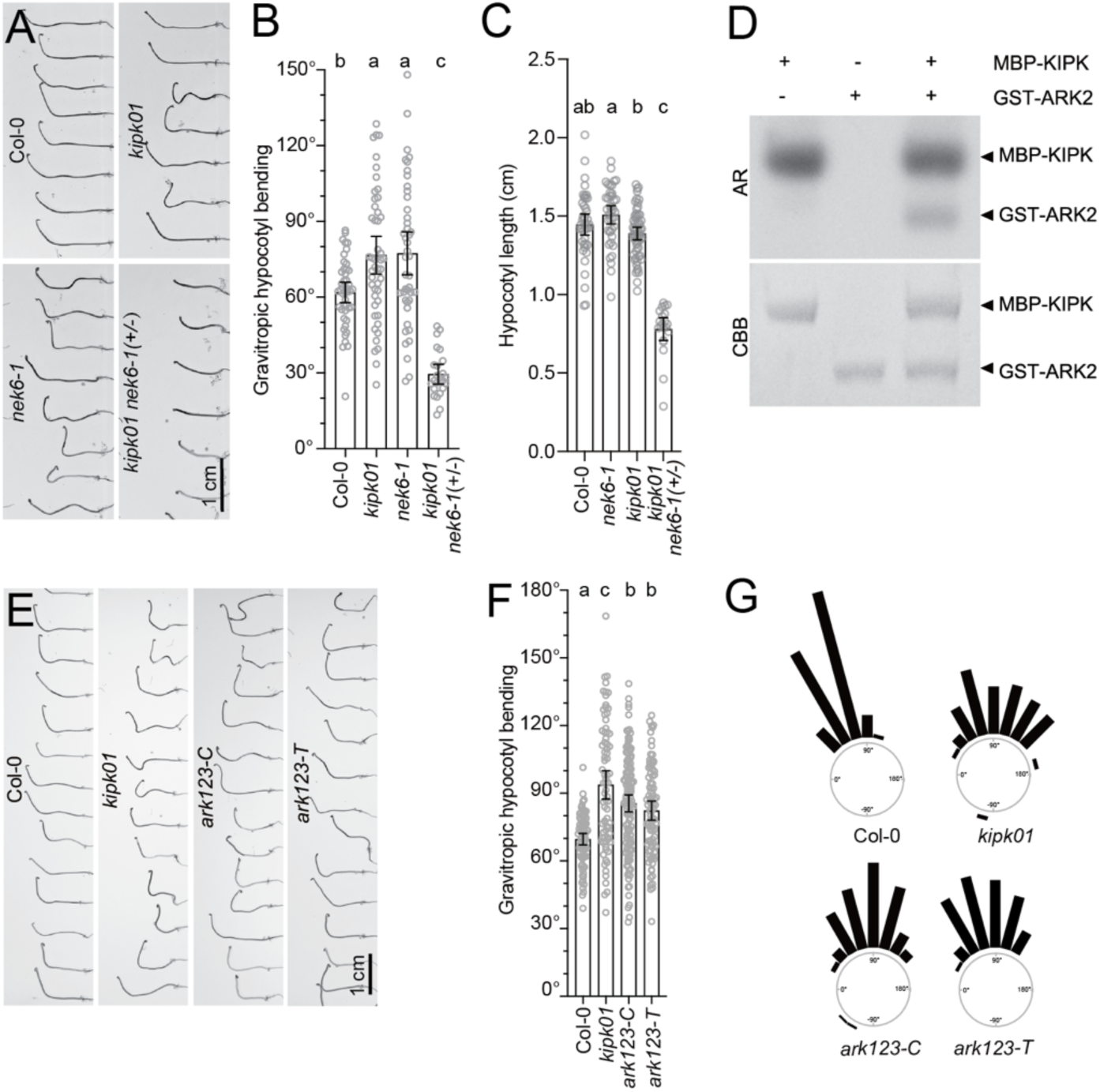
Interaction of KIPK with the ARK-regulatory kinase NEK6 and ARK kinesins. **(A)** and **(E)** Representative photographs of three-days-old dark-grown seedlings of the specified genotypes 24 hours after reorientation by 90°. **(B)** and **(F)** Graphs displaying the average and 95% confidence interval, as well as the individual data points from a negative hypocotyl gravitropism experiment as shown in (A) and (E), respectively. Results from a One-way ANOVA analysis are displayed on the top of each bar. **(C)** Graphs displaying the average and 95% confidence interval, as well as the individual data points from hypocotyl length measurements. Results from a One-way ANOVA analysis are displayed on the top of each bar. **(D)** Autoradiograph (AR) and Coomassie Brilliant Blue- (CBB-)stained gel of an SDS-PAGE with the result of *in vitro* phosphorylation reactions with MBP-KIPK and GST-ARK2. **(G)** Rose diagrams displaying hypocotyl bending angles of seedlings from the experiments shown in (E) and (F). Seedlings were grouped in 15° angle windows. n > 79 seedlings.

### KIPK phosphorylates ARK kinesins

The *NEK6* gene dosage-dependent synergistic interaction between *KIPK*, *KIPKL1* and *NEK6* suggested that the two types of kinases may have redundant biochemical function, e.g. in the regulation of the kinesin ARK2 that had previously been identified as NEK6 target (Sakai et al., 2008). When we tested recombinant purified GST-tagged ARK2 in combination with MBP-KIPK, we detected phosphorylation of ARK2 by KIPK, suggesting that ARK2 is a phosphorylation target of KIPK (Figure 8D).

ARK2 is highly homologous to ARK1 and ARK3, which may have redundant function (Sakai et al., 2008). We found that the T-DNA insertion in the 5’-untranslated region of *ARK3* in a previously characterized *ark1 ark2 ark3* (*ark123-T*) triple mutant still expressed high levels of *ARK3* and concluded that it may not be a loss-of-function mutant (Figure S10). We therefore generated a new *ark1 ark2 ark3* (*ark123*-C) triple mutant using CRISPR/Cas9 mutagenesis (Supplementary Figure S10C). When we examined *ark123*-C and *ark123-T* for gravitropic hypocotyl bending, we found that both mutant combinations displayed a similar bending defect as *kipk01* (Figure 8E and F). We concluded that KIPK, and most likely also KIPKL1, act redundantly with NEK6 kinases and that both types of kinases may exert their function during hypocotyl bending through the regulation of ARK kinesins.

### *ZWI* and *PERK8 PERK9 PERK10* are not required for gravitropic hypocotyl bending

The interaction of KIPK with kinesins of the ARK family was interesting with regard to the original identification of KIPK as an interactor of KINESIN-LIKE CALMODULIN-BINDING PROTEIN (KCBP), which had also been identified based on the trichome formation mutant *zwichel* (*zwi*) (Oppenheimer et al., 1997; Day et al., 2000). When we examined hypocotyl gravitropism in the *zwi* mutant, we did, however, not observe negative gravitropic bending defects in *zwi* mutants (Supplementary Figure S11A and B). Similarly did we not observe a corresponding phenotype in mutants of the previously identified PERK family KIPK interactors when examining *perk8 perk9 perk10* triple mutants (Supplementary Figure S11C and D) (Humphrey et al., 2015). We therefore concluded that these two interactors and their homologues may not act together with KIPK and KIPKL1 in the context of gravitropic hypocotyl bending.

### *KIPK* and *KIPKL1* are required for efficient seedling penetration through the soil

Negative gravitropic growth of elongating hypocotyls is essential for soil penetration during etiolated seedling growth after seed germination. However, in the soil and with the limited resources of the seed, elongating hypocotyls constantly need to adapt their growth direction to efficiently grow around obstacles and resume negative gravitropic growth to efficiently reach the sunlight for autotrophic growth. We reasoned that KIPK and KIPKL1 may be essential for efficient soil penetration of germinated seedlings. To test this hypothesis, we buried wild type and *kipk01* mutant seed 1 cm under the soil surface and, as a control, germinated the seedlings on the soil surface. While approximately 75% of the wild type seedlings emerged from the soil under these conditions, this number was reduced to 50% in the case of *kipk01* mutants (Figure 9A and B). In both populations, *kipk01* seedlings that had emerged from the soil and *kipk01* seedlings that had not penetrated from the soil, we observed enhanced hypocotyl bending and curving, which we quantified by determining seedling straightness (Figure 9C and D). Naturally, these phenotypes were more pronounced in seedlings that had not penetrated from the soil (Figure 9C and D). We concluded that KIPK and KIPKL1 are required for re-establishing negative gravitropic growth of seedling hypocotyls after their reorientation forced by contact with soil particles.

**Figure 9.**
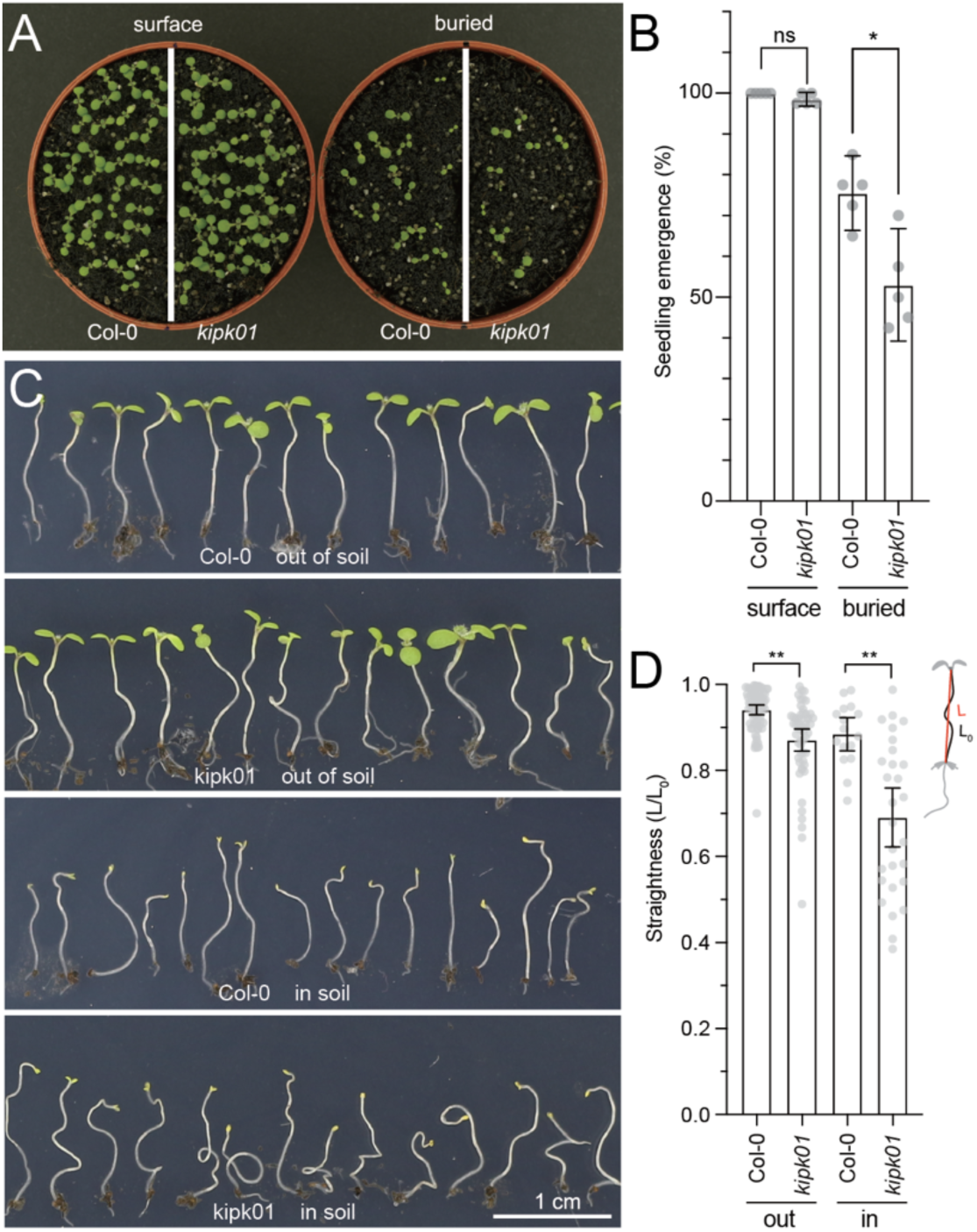
*KIPK* and *KIPKL1* are required for efficient soil penetration of seedling hypocotyls. **(A)** Top view of wild type (Col0) and *kipk kipkl1* mutants seedlings sawn on the surface of soil or buried 1 cm below the soil. **(B)** Graph displaying the average and 95% confidence interval, as well as the individual data points from experiments with seeds sawn on the soil surface or burried in the soil. **(C)** Photographs of seedlings germinated in the soil after emergence from the soil (out of soil) or without emergence from the soil (in soil). Scale bar = 1 cm. **(D)** Graph displaying the average and 95% confidence interval, as well as the individual data points from straightness measurements of seedlings germinated in the soil after emergence from the soil (out) or without emergence from the soil (in), as determined by division of the lengths (L_0_) and the heights (L) of the seedling hypocotyls. Student’s t-test. * p < 0.05, ** p < 0.01, *** p < 0.001, ns not significant.

## DISCUSSION

In the present study, we identify KIPK and KIPKL1 as redundant regulators of proper hypocotyl bending during negative gravitropic responses. The overbending phenotypes can be observed in *kipk kipkl1* mutants after being exposed to a change in the gravitropic vector, as well as in *kipk kipkl1* mutants growing through soil where the soil particles represent growth restricting obstacles that mechanically induce changes in the gravity vector. Our work identifies auxin transport regulation, as well as regulation of the ARK kinesins and hence cortical microtubules as two, mutually non-exclusive mechanisms of how KIPK and KIPKL1 control hypocotyl overbending in the wild type.

KIPK and KIPKL1 are polarly localized plasma membrane associated proteins whose membrane association is impaired in the presence of PAO (phenylarsine oxide), an inhibitor of phosphoinositol-4-phosphate synthesis (Simon et al., 2016). Although it is therefore very likely that the interactions at the plasma membrane, e.g. with phosphoinositol-4-phosphate or its derivatives, are mediated by interactions with positively charged basic amino acids in KIPK and KIPKL1, the mutation of a strongly polybasic motif with 12 basic amino acids found in the KIPK N-terminus did not abolish its plasma membrane association and did not impair the biological functionality of the protein in complementation experiments.

Although KIPKL2, the third family member, is highly homologous to KIPK and KIPKL1 and shares the pronounced polar distribution at the basal plasma membrane with KIPK and KIPKL1, our data clearly indicate that KIPKL2 has distinct functions. First, expression of *KIPKL2*, unlike the expression of *KIPK* or *KIPKL1* from a *KIPK* promoter fragment, cannot suppress the phenotype of the *kipk012* mutant. Second, the loss of *KIPKL2* in the *kipk012* triple mutant does not quantitatively enhance the phenotype of the *kipk01* double mutant. Third, in auxin transport assays, KIPKL2 is less efficient than KIPK and KIPKL1 in activating the auxin transporter PIN3. Then, while KIPKL2 shares the strong polar distribution at the basal plasma membrane with KIPK and KIPKL1 and their affinity for phospholipids, KIPKL2 differs from the other two family members with regard to the presence of a polybasic motif in the middle domain of KIPKL2, which is reminiscent of similar polybasic motifs in the middle domains of D6PK, D6PK-LIKE proteins or PAX but absent from KIPK or KIPKL1 (Barbosa et al., 2016; Bassukas et al., 2022). We conclude from these observations that KIPKL2 is neither biochemically nor biologically redundant with KIPK and KIPKL1. With regard to the latter point, we noted with interest that the promoter activity of *KIPKL2* in the meristematic zone of the root tip is suggestive for a cell cycle-dependent regulation. Further research is thus needed to elucidate the function of KIPKL2 and its functional distinction from KIPK and KIPKL1 at the biological, biochemical and cell biological level.

Like D6PK, D6PKL and PAX kinases, KIPK and KIPKL1 are polarly localized at the basal (rootward) plasma membrane, where they colocalize with polarly localized PINs, and phosphorylate ‘canonical’ PIN transporters *in vitro*. The requirement of PINs and auxin transport for the initiation of tropic bending responses, prevented us from examining their role during the termination of the bending response, which is impaired in the *kipk kipkl1* mutants or bending during hypocotyl elongation growth in the soil. Since we measured reduced auxin transport in *kipk kipkl1* mutant hypocotyls while auxin levels remained similar between the wild type and the *kipk kipkl1* mutants, we argue that a reduction of KIPK and KIPKL1-dependent PIN-mediated polar auxin transport, at least in part, contributes to the growth defects of *kipk kipkl1* mutants.

Previous work on hypocotyl gravitropic bending had suggested that the bending process was accompanied by a redistribution of the auxin transporter PIN3 in gravity-sensing endodermis cells, in line with the assumption that a relative increase in auxin distribution at the lower hypocotyl cell files would promote the cell elongation required for bending (Rakusova et al., 2011). Further, it had been postulated that the consequent auxin accumulation at the lower hypocotyl cell files would induce a second lateral redistribution of PIN3 to terminate the bending response (Rakusova et al., 2016; Grones et al., 2018). We reasoned that differences in this PIN3 distribution may be causal for the bending phenotypes oberved in *kipk01* mutants. However, when we examined the distribution of PIN3-GFP expressed from the endodermis-specific *SCR* promoter in cells undergoing differential elongation, we failed to observe the previously reported PIN3 redistributions between the two lateral membranes. Our observation that there is no dynamic redistribution of PIN3 between the lateral plasma membranes during gravitropic responses is in conflict to previously published work that described such a redistribution and also the termination of the bending response as a consequence of cellular auxin accumulation (Rakusova et al., 2011; Rakusova et al., 2016; Grones et al., 2018). From this point of view, we conclude that the previously established model of an auxin-regulated differential lateral PIN3 distribution during negative hypocotyl bending, although attractive, may require a further detailed re-examination. The absence of a PIN3-GFP redistribution also prevented us from using this cell biological readout for a further characterization of the overbending phenotype of *kipk kipkl1* mutants.

Another possible mechanism for the regulation of gravitropic hypocotyl bending through the polarly localized KIPK and KIPKL1 kinases would have been a cellular redistribution of the KIPK and KIPKL1 proteins during gravitropic response. However, similarly to our inability of observing such polarity changes with PIN3, we also did not observe changes in the polar distribution of KIPK or KIPKL1 during gravitropic responses when examined in the endodermis, a tissue file where *KIPK* expression from the *SCR* was sufficient to suppress the *kipk01* phenotype.

Based on the phenotypical similarities, we also considered a functional interplay between KIPK, KIPKL1 and ABC family auxin transporters. We found, however, that mutant of the *ABCB19* transporter had different defects in auxin transport than *kipk kipkl1* mutants with regard to the defects in auxin transport observed in *kipk kipkl1* mutants and their sensitivity to the inhibitor NPA. We therefore considered a functional relationship between the kinases and the ABCB transporters unlikely. Further, recent work has shown that brassinosteroids rather than auxins may be the primary cargoes of the ABCB transporters (Ying et al., 2024).

Our research identifies KIPK interactions with members of the five-membered BRX/BRXL protein family. We identify BRXL2 as a KIPK phosphorylation target and show that BRXL2 and BRXL4 share the basal polar localization with KIPK and KIPKL1. We characterize BRXL2 S292 as a dominant KIPK-targeted phosphorylation site and find that the site is conserved between BRXL2, BRLX3, and BRXL4, which form a subfamily among the BRX/BRXL protein family. In previous work, we had linked BRX, the founding member of the family, with the regulation of the AGC1 kinase PAX in the context of protophloem differentiation (Marhava et al., 2018; Koh et al., 2021). There, we had observed repressive effects of BRX on PAX-mediated PIN activation when BRX and PAX are co-expressed. At the same time, BRX had been reported to be an auxin-labile polar plasma membrane protein, which, together with other findings, led to the conclusion that BRX was part of an auxin-regulated mechanism where BRX would repress PAX and consequently auxin transport at low cellular auxin concentrations (Marhava et al., 2018; Koh et al., 2021). This repression would be relieved through BRX degradation at elevated auxin levels leading to a PAX-regulated PIN-dependent auxin efflux (Marhava et al., 2018). Further work had shown that BRXL2 was functionally distinct from BRX (Koh et al., 2021). Here, we find that BRXL2 does not have an effect on KIPK- or KIPKL-dependent PIN activation in oocytes. Still, the possibility of a dynamic regulation of auxin transport during gravitropic bending or obstacle avoidance by BRXL2 or BRXL2 phosphoregulation by KIPK and likely also KIPKL1 is very intriguing since it would pose the possibility for the dynamic growth regulation of this auxin- and auxin transport-dependent process. Further research is needed to examine this proposed interplay at the cell biological and biochemical level. With regard to our observation of BRXL proteins having a role in tropic responses, we would like to emphasize that BRXL proteins, as well as RLD (RCC1-LIKE DOMAIN) proteins, which contain a so-called BRX domain while being distinct from the *bona fide* BRX/BRXL protein family, have been implicated in the regulation of tropic responses in conjunction with LAZY family proteins (Li et al., 2019; Furutani et al., 2020; Waite et al., 2024). LAZY proteins, in turn, were recently found to be transferred from the amyloplast membrane to the interacting plasma membrane after amyloplast sedimentation following a change in the gravitropic vector (Furutani et al., 2020; Chen et al., 2023; Kulich et al., 2023; Nishimura et al., 2023). One of these recent studies even described the recruitment of the AGC1 kinase D6PK to LAZY at the plasma membrane (Kulich et al., 2023). While root gravitropic responses are not impaired in the *kipk kipkl1* mutants, the relatedness of LAZY and BRX/BRLX proteins may invite the hypothesis that the two types of proteins function in a cell biologically or biochemically similar manner in the recruitment or regulation of AGC1 family kinases for localized responses, such as localized auxin transport or localized regulation of microtubules. Dedicated studies of auxin distribution and auxin transporter activities, on the one side, and microtubule distribution and activity, on the other side, will be required to examine this interaction and their effects in the future.

Besides BRXL family proteins, we also identify ARK kinesins are being required for proper gravitropic responses and hypocotyl bending. The link between ARK kinesins, on the one side, and KIPK and KIPKL1, on the other, was made because mutants of the ARK regulatory kinase *NEK6* displayed a hypocotyl overbending phenotype reminiscent of the phenotype observed in *kipk01* mutants. We subsequently observed a strong synergistic interaction between *kipk01* and *nek6* mutants and that KIPK, like the structurally unrelated NEK6 kinase, can phosphorylate ARK kinesins *in vitro*. The fact that *kipk01*, *nek6* and *ark123* triple mutants display similar phenotypes suggests that these proteins act, indeed, in the same pathway and control gravitropic bending responses. The strong genetic interaction between KIPK, KIPKL and NEK kinases also suggests that KIPK and KIPKL1 have other biological functions, besides the function in gravitropic bending reported here, that may be genetically buffered in other biological contexts by *NEK6* or other, as-yet unknown regulators.

NEK6 has been implicated in the regulation of the microtubule network during cellular tensile stress as it occurs during gravitropic bending or other uneven bending responses (Takatani et al., 2020). Similarly, the proposed regulation of ARK kinesins by KIPK strongly supports the notion that, besides PIN-mediated auxin transport regulation, the regulation of cortical microtubule dynamics may be a cellular mode of growth regulation by KIPK kinases. It can also be easily imagined that the regulation of microtubule dynamics is a mode of action whereby other members of the AGC kinase family regulate plant growth and development at the cell biological level, e.g. during phototropic hypocotyl bending, which is mediated by D6PK and D6PKL AGC1 kinases or the phototropin AGC4 kinases. The analysis of microtubule networks during gravitropic bending in the wild type and the *kipk kipkl1* mutants would require a dedicated effort that goes beyond the scope of the present study.

Our observation of a KIPK/KIPKL interaction with ARK kinesins is also interesting in the context of the original characterization of KIPK as interactor of the kinesin ZWI (Day et al., 2000). Although *kipk kipkl1* did not display reductions in trichome branch numbers as reported for *zwi* and, inversely, *zwi* mutants did not display the overbending phenotype of *kipk kipkl1* mutants, there may be, as yet unknown, biological processes where a ZWI kinesin regulation by KIPK/KIPKL kinases may be biologically relevant. Similarly, *kipk kipkl1* hypocotyl bending phenotypes cannot be observed in mutants of the reported KIPK-interacting *PERK* kinases (Humphrey et al., 2015). PERK kinases are cell surface proteins with extension motifs and cytoplasmic kinase domains (Borassi et al., 2016). PERKs have a proposed role in cell wall integrity sensing that link cell wall interactions through repeated extension motifs with cytoplasmic signalling (Borassi et al., 2016). In line with this proposed function, gravitropic bending involves cell expansion, requiring cell wall loosening, deposition of new cell wall materials, and subsequent cell wall rigidification and, thus, could require proteins with the proposed functionality of PERKs. Thus, the reported interaction between KIPK, KIPKL1 and PERKs may deserve further research and could reveal biological functions beyond the reported role of PERKs in root growth on media containing high sucrose levels.

In summary, our study uncovers a previously unknown biological role of KIPK and KIPKL1 proteins in the regulation of gravitropic hypocotyl bending that is biologically also relevant during the penetration of seedling hypocotyls through soil. We provide evidence that the regulation of PIN-mediated auxin transport may be one mechanism whereby KIPK and KIPKL1 regulate this tropic growth response, but also uncover a role in ARK kinesin regulation as a previously unknown function of these kinases. While it can easily be envisioned that these findings can also radiate to the role of other AGC kinase family members, the strong genetic interaction with NEK6 further highlights that there may be other biological functions of KIPK and KIPKL1 that remain to be discovered.

## METHODS

### Biological material

All experiments were performed with *Arabidopsis thaliana* Columbia-0 (Col-0) as wild type control. The insertion alleles *kipk-1* (SALK_065651C), *kipkl1-1* (GABI_175F10), *kipkl1-2* (SALK_200701C), *kipkl2-1* (SALK_087379C) and *kipkl2-2* (SALK_015563) were obtained from the Nottingham Arabidopsis Stock Center (NASC, Nottingham, UK) and genotyped for homozygosity before use for experimental work. The alleles *kipk-1*, *kipkl1-1* and *kipkl2-1* were used to generate *kipk01*, *kipk02*, *kipk12* double and *kipk012* triple mutants. *kipk-1* and *kipkl1-2* were used to generate the *kipk01-2* double mutant.

The following mutants and alleles have been described elsewhere: *ark1-1 ark2-1 ark3-1* (Sakai et al., 2008); *abcb19* (SALK_033455) (Lewis et al., 2007); *brxl2* (SALK_032250) and *brxl3* (SALK_017909) (Briggs et al., 2006); *brxl4-1* (SALK_147349c) and *brxl4-2* (SALK_022411c) (Che et al., 2022)(Che et al., 2023); *35S::YFP-D6PK* (Zourelidou et al., 2009); *d6pk d6pkl1 d6pkl2* (*d6pk012*) (Zourelidou et al., 2009); *nek6-1 (Motose et al., 2008)*; *pin3-3* (2 bp deletion) (Friml et al., 2002); *pin3-3 pin4-101* (GABI_593F01) *pin7-102* (SALK_062056) (*pin347*) (Willige et al., 2013); pSCR::PIN3-YFP (Rakusova et al., 2011); pPIN3::PIN3-GFP (Zadnikova et al., 2010); *zwi* (SALK_031704) (Humphrey et al., 2015); *perk8-1* (SALK_129961) *perk9-1* (SALK_014687) *perk10-1* (SALK_022872) (Humphrey et al., 2015).

Primers used for genotyping are listed in Supplementary Table S1.

### Plant cultivation

For growth in sterile culture, seeds were surface-sterilized for seven minutes in 25% DanKlorix hygiene cleaner (CP GABA, Hamburg, Germany) and 0.05% Triton X-100, followed by five washes with sterile water. Sterilized seeds were kept in the dark at 4°C for two to four days for stratification and germinated on ½ Murashige and Skoog (½ MS) medium (Duchefa, Harlem, The Netherlands) supplemented with 0.05% 2-(N-morpholino) ethanesulfonic acid, 1% sucrose, and 0.6% or 0.8% agar, depending on whether the seeds were grown on horizontally (0.6%) or vertically (0.8%) oriented plates. Seedlings on plates were grown at 21°C in the dark or in continuous light (110 µM m^−2^ s^−1^). For growth on soil, plants were grown at 21°C with 16 h photoperiod (120 µM m^−2^ s^−1^) and 65% relative humidity.

### Cloning procedures and transgenic lines

pGreenII0229-pKIPK::YFP-KIPK (pKIPK::YFP-KIPK), expressing a genomic variant of *KIPK*, was used for phenotype complementation. To obtain pGreenII0229-pKIPK::YFP-KIPK, a genomic fragment of *KIPK* was introduced into pDONR201™ and subcloned into pGreenII0229-pD6PK::YFP-GW using Gateway methodology to obtain pGreenII0229-pD6PK::YFP-KIPK (Invitrogen, Carlsbad, CA). Subsequently, the *D6PK* promoter fragment was replaced by a 2 kb *KIPK* promoter fragment flanked by *KpnI* and *XhoI* sites in pGreenII0229-pD6PK::YFP-KIPK.

To generate constructs for *KIPK* or *KIPKL* overexpression, their full length coding sequences were PCR-amplified from Col-0 cDNA and cloned into pDONR201™ using BP Clonase (Invitrogen, Carlsbad, CA) and subsequently subcloned into pUBN-Dest-eGFP-GW (Grefen et al., 2010) to obtain pUBN::eGFP-KIPK (eGFP-KIPK), pUBN::eGFP-KIPKL1 (eGFP-KIPKL1) and pUBN::eGFP-KIPKL2 (eGFP-KIPKL2).

The *UBN10* promoter fragment between the *EcoRI* and *XhoI* sites was then replaced by 2 kb *KIPK* or *KIPKL1* native promoter fragments, to obtain, using standard cloning procedures, pKIPK::eGFP-KIPK and pKIPKL1::eGFP-KIPKL1 for mutant complementation. *KIPKL2*, *D6PK* and *KIPK* mutant variants were introduced by Gateway LR cloning (Invitrogen, Carlsbad, CA) into pKIPK::eGFP-GW, which had been obtained with a BP Clonase (Invitrogen, Carlsbad, CA) recation between pKIPK::eGFP-KIPK and pDONR201, to obtain pKIPK::eGFP-KIPKL2, pKIPK::eGFP-D6PK and pKIPK::eGFP-KIPK mutant variants.

For endodermis-specific expression of *KIPK*, a 2 kb *SCARECROW* (*SCR*) promoter fragment and a *BsaI* site-cured *KIPK* coding sequence were ligated into pGGA000 and pGGC000 entry vectors, respectively, to subsequently generate pFASTR-pSCR::mCitrine-KIPK (pSCR::mCitrine-KIPK) in pFASTR-AG by Greengate cloning (Lampropoulos et al., 2013; Decaestecker et al., 2019).

Point mutations in *KIPK* were introduced by site-directed mutagenesis into pDONR201-KIPK as a template to obtain the kinase-dead K567E variant and the 12RKA variant with mutations of 12 basic amino acids defining the domain with a high (> 0.6) BH score located between R332 and K355 (R332A, K333A, R336A, R340A; K342A, K345A, K346A, K347A, K351A, K352A, K353A, K355A) (Fisher and Pei, 1997).

To generate pKIPK::eGFP-GUS, as well as KIPKL::eGFP-GUS vectors, 2.0 kb *KIPK* and *KIPKL* promoter fragments were amplified from Col-0 genomic DNA, cloned into pDONR201™ and subsequently subcloned into pFASTG04 using Gateway technology (Invitrogen, Carlsbad, CA) (Shimada et al., 2010).

All constructs for the expression of *KIPK/KIPKL* or mutant variants under the control of the *pKIPK*, *pKIPKL1*, *pKIPKL2* or *pSCR* promoters were transformed into the *kipk01* mutant to analyse mutant complementation and protein localization. pKIPK/pKIPKL::eGFP-GUS constructs were transformed into Col-0 for gene expression pattern visualization. For each construct, homozygous lines of at least three independent lines were analysed.

To obtain modified DR5V2::GUSm, the previously published DR5V2::GUS vector (Hayes et al., 2019), lacking a terminator sequence for *ß-glucuronidase* (*GUS*) expression, was modified by introducing a 256 bp terminator fragment of *HEAT SHOCK PROTEIN 18.2* into the *SacI* site. DR5V2::GUSm was transformed into *kipk01* and back-crossed to the Col-0 wild type to analyse auxin distribution.

The coding sequences of *KIPK*, *KIPKL1*, *KIPKL2* were ligated into the blunt ended pOO2 vector, obtained by E*co32* I, S*ma* I, and E*co32* I restriction digestion, respectively.

To obtain CRISPR/Cas9 vectors for creating mutants of *BRXL* and *ARK* genes, two sgRNAs were designed for each gene. sgRNA1-U6_26t-U6_29p-sgRNA2 were amplified from the pCBC-DT1T2 vector by using the primers containing the sgRNA sequences and ligated into pENTR-MSR (Xing et al., 2014). Three U6_26p_sgRNA1-U6_26t-U6_29p-sgRNA2-U6_26t fragments for *BRXL2*, *BRXL3*, and *BRXL4*, as well as for *ARK2*, *ARK3*, and *ARK4* from pENTR-MSR were clustered by isocaudamer enzyme-ligation with SpeI and *Xba*I. The whole sgRNA cluster was subcloned into the *Spe*I-*Hind*III site of pHEE401E (Wang et al., 2015; He et al., 2022).

To obtain constructs for protein expression and purification in *Escherichia coli*, GST-fused *KIPK/KIPKL* and ARK2 were cloned by Gateway technology into pDEST15^TM^. MBP-fused *KIPK/KIPKL* and GST-fused BRXL2 variants were introduced by Greengate cloning into pGGP000, a gift from Andrea Bleckmann (University of Regensburg, Germany) (Lampropoulos et al., 2013).

All primers used for cloning and mutagenesis are listed in Supplementary Table S1.

### Protein alignment

Full length *Arabidopsis thaliana* AGC1 and AGC4 protein sequences were used for protein alignment with the ClustalW algorithm of the Geneious software (Thompson et al., 1994). Phylogenetic analysis was done using the MEGA-X software with the Neighbour-Joining method (Saitou and Nei, 1987).

### Semi quantitative RT-PCR

Total RNA was extracted from five-days-old light-grown seedlings using the NucleoSpin RNA Plant kit (Macherey Nagel, Düren, Germany). cDNA was synthesized with a RevertAid First Strand cDNA synthesis kit (Thermo Scientific, Waltham, MA) and the *KIPK/KIPKL* open reading frames were amplified with DreamTaq DNA Polymerase (40 cycles; Thermo Scientific, Waltham, MA). ACTIN2 was analysed as a control gene (20 cycles). Primers are listed in Supplementary Table S1.

### Tropism and soil growth assays

For hypocotyl negative gravitropism assays, seeds were first germinated for 18 – 24 hrs in the light and then grown for 2.5 – 3 days in the dark on vertically oriented plates covered in multiple layers of aluminium foil. The hypocotyls of the etiolated seedlings were straightened in th direction of the gravity vector in safe green light and allowed to recover for 1 - 2 hours. For negative hypocotyl gravitropism assays, the plates were turned by 90° for gravity stimulation. For phototropism assays, seedlings were prepared in an identical manner and plates were then transferred to a FloraLED chamber (CLF Plant Climatics, Wertingen, Germany) and illuminated with 1 μmol m^−2^ s^−1^ blue light from a 90° angle. After 24 hours (negative hypocotyl gravitropism) or 16 hours (phototropism). For testing the effect of light quality on *kipk01* phenotype, seedlings were illuminated with 0.05 μmol m^−2^ s^−1^ blue light or 0.1 μmol m^−2^ s^−1^ red light for 4 days in a FloraLED chamber (CLF Wertingen, Germany). All the hypocotyls were photographed with a Canon EOS 650D camera (Canon, Tokyo, Japan) and hypocotyl curvature was determined with ImageJ (Fiji) (https://imagej.nih.gov/ij/). Root positive gravitropism was measured from six-days-old light-grown seedlings that had been aligned in the direction of the gravity vector on vertically oriented plates. Roots were photographed eight hours after reorienting the plates by 90° and root bending was determined with ImageJ (Fiji) (https://imagej.nih.gov/ij/). For measurements of shoot negative gravitropism, *Arabidopsis thaliana* plants with ca. 10 cm inflorescence height were reoriented by 90° for gravity stimulation and photographed with a Canon EOS 750D camera (Canon, Tokyo, Japan) in 1 hr intervals. Shoot angles were measured using ImageJ (Fiji) (https://imagej.nih.gov/ij/). GraphPad Prism 9 (GraphPad Software, San Diego, CA) and Origin 8.0 (OriginLab, Northhampton, MA) were used for statistical analyses and plotting.

### GUS staining

To determine *KIPK/KIPKL* expression from GUS-GFP reporter lines or auxin distribution from DR5V2::GUS, three-days-old dark-grown seedlings or three- and five-days-old light-grown seedlings were fixed in 90% acetone at −20°C for 2 h, washed twice with water before placing the seedlings into GUS staining buffer (10 mM EDTA, 50 mM sodium phosphate [pH 7.0], 0.1% [v/v] Triton X-100, 0.5 mM K_3_Fe(CN)_6_, 0.5 mM K_4_Fe(CN)_6_, 1 mg/mL X-GlcA). Samples were placed under vacuum for 30 min to improve penetration of the staining solution. *proKIPK/KIPKL::GUS-GFP*, DR5V2::GUS and *DR5V2::GUS kipk01* were incubated at 37°C for 18, 4 and 24 hours respectively. 4 hour-staining did not show any visible signal in the hypocotyl of *DR5V2::GUS kipk01* mutant. All the samples were cleared in 70% (v/v) ethanol, and imaged with a Leica MZ16 microscope equipped with a Leica DMC5400 camera system (Leica, Wetzlar, Germany).

To determine seedling growth in soil, Arabidopsis seeds were sown on 1/2 MS agar plates and exposed to light for 6 hrs. Absolute seed number was determined and seeds were transferred to the soil surface or buried under 1 cm of loamy soil, prepared by blending sieved organic matter with sand (average diameter 1mm) in a 1:5 ratio based on dry weight. Seed germination and seedling emergence were assessed after growth at 22°C for 7 days.

### Confocal microscopy

To visualize YFP- and eGFP-fusions of KIPK and D6PK AGC kinases in hypocotyls and roots, respective seedlings were cleared with ClearSee or ClearSeeAlpha for two to four days (Kurihara et al., 2015; Kurihara et al., 2021). pSCR::PIN3-YFP and pPIN3::PIN3-GFP were directly detected in three-days-old dark-grown seedlings at the specified time points with an Olympus FV1000 confocal microscope with a high-sensitivity detector unit using a 515 nm laser and a 520 - 550 nm band pass filter for YFP detection and a 488 nm laser and a 505 – 540 nm band pass filter for eGFP detection (Olympus, Tokyo, Japan). The mean intensities of PIN3-YFP on the inner and outer side of the endodermis were measured with ImageJ (Fiji) by drawing a free-hand line along the entire lateral plasma membrane at each of the two lateral sides of the cell (https://imagej.nih.gov/ij/).

### Chemical inhibitor experiments

For chemical inhibitor treatments, roots of five-days-old light-grown seedlings expressing YFP-KIPK and YFP-D6PK were imaged; the respective concentration and treatment duration for each chemical is indicated in figure legends as previously described (Gao et al., 2013; Barbosa et al., 2016). Taxol and Isoxaben (Sigma, Taufkirchen, Germany) were directly added into ½ MS medium at the specified concentrations; seedlings were grown for four days in the dark on vertically oriented plates covered in multiple layers of aluminium foil prior to imaging and hypocotyl length measurements with ImageJ (Fiji) (https://imagej.nih.gov/ij/).

### PIP strip experiment

For recombinant protein expression and purification, GST-KIPK and GST-KIPKL proteins were expressed in *Escherichia coli* Rosetta (DE3). Free GST protein was expressed from a pGEX-KG empty vector as a negative control (ATCC, Manassas, VA) (Guan and Dixon, 1991). All GST-tag proteins were purified using Protino Glutathione Agarose 4B (Macherey-Nagel, Düren, Germany). Lipid binding assays were performed following the previous description using TBS buffer on PIP-strips P-6001 (Echelon, San José, CA) (Barbosa et al., 2016).

### *In vitro* phosphorylation experiments

For *in vitro* phosphorylation reactions, MBP fusion proteins were expressed in *Escherichia coli* Rosetta (DE3) pLysS (Novagen) and purified using Amylose Resin (New England Biolabs, Ipswich, MA). 0.5 - 1 μg purified GST-PIN3 cytoplasmic loop (CL) and 0.1 - 0.2 µg recombinant kinase were used for each reaction in 1x kinase buffer (25 mM Tris pH 7.5, 5 mM MgCl_2_, 1 mM DTT), 100 μM ATP and 3 μCi [γ-^32^P] ATP [10 μCi μl^−1^ stock] (Hartmann Analytic, Braunschweig, Germany). Reactions were incubated at 30°C for 1 hour and stopped by addition of 5x Laemmli SDS loading buffer and boiling at 56°C for 10 min. Reactions were subsequently separated on a 4-12 % iD PAGE Gel (Eurogentec, Liege, Belgium). The gels were washed twice with 5% trichoroacetic acid, stained with Coomassie Brilliant Blue (CBB), destained with destaining buffer (40% ethanol, 7% acetic acid) and dried overnight. Autoradiography was performed for six hours (MBP-KIPK) or five days (MPB-KIPKL1 and MBP-KIPKL2) at −80°C on X-ray film (CEA, Cadarache, France).

### *In vivo* auxin transport assay

Auxin transport assays in hypocotyls of etiolated seedlings were carried out as previously reported, with slight modifications (Willige et al., 2013). Briefly, ^3^H-IAA (25 Ci mmol^−1^; RC Tritec, Teufen, SWITZERLAND) was dissolved to a final concentration of 400 nM in 5 mM MES (pH 5.5), 1% glycerol. Four-days-old dark-grown seedlings were transferred onto Parafilm (Beemis Company, Neenah, WI) strips placed on the surface of ½MS vertical plates. Seedlings were aligned on a 6 mm Parafilm strip with the cotyledons and the apical part of the hypocotyl covering 5 mm of the parafilm strip, allowing for the application of 0.5 μl ^3^H-IAA solution to the cotyledons of each seedling. Seedlings were then incubated vertically in the dark for four hours. Subsequently, the lower part of the hypocotyl was excized. For scintillation counting, five seedlings were grouped into one scintillation vial containing 2 ml Ultima Gold liquid scintillation cocktail and counted in a Tri-Carb 4910TR Liquid Scintillation counter (Perkin Elmer, Rodgau, Germany).

### Oocyte auxin efflux assay

For auxin transport assays conducted in oocytes, *Xenopus laevis* oocytes were collected as previously described (Zourelidou et al., 2014; Fastner et al., 2017). cRNA was synthesized using the mMessage Machine SP6 Kit (Life Technologies, Carlsbad, CA) and cRNA concentration was adjusted to 300 ng/µl PIN and 150 ng/µl protein kinase, respectively. Oocytes were injected the day after surgery with ∼50 nl of a 1:1 mixture of cRNAs for PIN and the respective protein kinase. If only PIN or protein kinase cRNA was injected, the cRNA was mixed 1:1 with water (mock control). Following injection, oocytes were incubated in Barth’s solution containing 88 mM NaCl, 1 mM KCl, 0.8 mM MgSO4, 0.4 mM CaCl2, 0.3 mM Ca(NO3)2, 2.4 mM NaHCO3, 10 mM HEPES (pH 7.4) supplemented with 50 µl gentamycine at 16°C for four days to allow for protein synthesis. An outside medium buffer at pH 7.4 was chosen to minimize passive rediffusion of IAA into the oocytes, which increases at acidic pH. At the beginning of the experiment, ten oocytes per time point were injected with 50 nl of a 1:5 dilution (in Barth’s solution) of [^3^H]-IAA, 25 Ci/mmol; 1 mCi/ml (ARC, St. Louis, MO or Tritec, Teufen, SWITZERLAND) to reach an intracellular oocyte concentration of ∼1 µM [^3^H]-IAA based on an estimated oocyte volume of 400 nl (Broer, 2010). After [^3^H]-IAA injection, oocytes were placed in ice-cold Barth’s solution for two mins to allow substrate diffusion and closure of the injection spot. Subsequently, oocytes were washed and transferred to Barth’s solution at 21°C to allow for auxin efflux. To stop auxin efflux, oocytes were washed twice and lysed individually in 100 µl 10% SDS (w/v) at selected time points and the residual amount of [^3^H]-IAA in each oocyte was determined by liquid scintillation counting. At least seven oocytes were measured per time point and mock as well as other negative controls were performed with the same oocyte batch to account for differences between batches. The relative transport rates of an experiment were determined by linear regression. Transport rates of different biological replicates, i.e. oocytes collected from different donor animals, were averaged and are presented as mean and standard error of at least three biological replicates.

### Auxin measurements

IAA measurements were performed from hypocotyls and cotyledons of four-days-old dark-grown seedlings as previously described (Simura et al., 2018).

### Immunoprecipitation and immunoblot analyses

Three-days-old dark-grown seedlings of *pPIN3::PIN3-GFP* in Col-0 and *kipk01* background were turned 90° and kept for two hours as gravity-stimulated samples. Gravity-stimulated seedlings and non-stimulated control seedlings were ground in liquid nitrogen and homogenized in extraction buffer (50 mM Tris-HCl [pH 7.5], 150 mM NaCl, 0.2% Triton X-100, 100 μM MG132, 1 mM PMSF, 1x cOmplete Protease Inhibitor Cocktail (Roche, Penzberg, Germany), and 1x PhosSTOP (Roche, Penzberg, Germany). Extracts were cleared by centrifugation at 5,000 *g* for five min. The supernatant was incubated with GFP-Trap Magnetic Agarose (Chromotek, Planegg, Germany) at 4°C for two hours. The beads were washed three times with extraction buffer. 400 U Lambda Protein Phosphatase (New England Biolabs, Frankfurt, Germany) in NEBuffer, containing 10 mM MnCl_2_, and purified GST-KIPK in kinase buffer (25 mM Tris pH 7.5, 5mM MgCl_2_, 1 mM DTT, 1 mM ATP) were added to two of the PIN3-GFP immunoprecipitates, respectively. Dephosphorylation and phosphorylation reactions were both incubated at 30°C for one hour. All protein samples were denatured at 42°C for ten min, after adding 5x Laemmli buffer. An anti-GFP (laboratory stock) and an anti-rabbit HRP-conjugated secondary antibody (Sigma, Taufkirchen, Germany) were used for western blot detection. Chemiluminescence was generated with SuperSignal West Femto Maximum Sensitivity Substrate (Thermo Scientific, 34096) and detected with a Fujifilm LAS 4000 mini (Fuji, Tokyo, Japan). Coomassie Brilliant Blue-stained gels of the lysate after homogenization were used as loading control.

### Co-immunoprecipitation and mass spectrometry

For co-immunoprecipitation-mass spectrometry, four-days-old dark-grown seedlings were harvested under safe green light and subsequently ground in extraction buffer. Supernatants were incubated with anti-GFP beads (GFP-Trap_MA; Chromotek, Martinsried, Germany). Beads were magnetically separated and protein was eluted in 2 × SDS sample buffer, loaded on an SDS–PAGE gel for electrophoresis, and subsequently analysed by liquid chromatography with tandem mass spectrometry. For tryptic digestions, samples were reduced and alkylated by 50 mM DTT and 10 mg/mL chloroacetamide, respectively. Tryptic in-gel digestion was performed according to standard procedures. Nano-flow liquid chromatography tandem mass spectrometry was performed on an Ultimate 3000 UHPLC system (Thermo Fisher Scientific) coupled online to an Orbitrap Eclipse Tribrid mass spectrometer (Thermo Fisher Scientific). Briefly, peptides were delivered to a trap column (100 μm × 2 cm, packed in house with Reprosil-Pur C18-AQ 5 µm resin; Dr. Maisch) at a flow rate of 5 µL/min in 100% solvent A (0.1% formic acid in HPLC-grade water). After 10 min of loading and washing, peptides were transferred to an analytical column (75 µm × 40 cm, packed in house with Reprosil-Pur C18-GOLD, 3 µm resin; Dr. Maisch) and separated using a 60-min gradient from 4 to 32% of solvent B (0.1% formic acid and 5% DMSO in acetonitrile; solvent A: 0.1% formic acid and 5% DMSO in water) at 300 nL/min flow rate. Tandem mass spectra were acquired in DDA mode. MaxQuant (version 1.6.17.0) with its built-in search engine Andromeda was used for peptide and protein identification and quantifications. MSMS spectra were searched against the Araport database (Araport11_genes.201606.pep.fasta; downloaded June 2016). Unless otherwise specified, the default parameters of MaxQuant were used. Precursor tolerance was set to ±4.5 ppm and fragment ion tolerance to ±20 ppm (FTMS) and 0.4 Da (ITMS). Trypsin/P was chosen as the proteolytic enzyme with up to two missed cleavage sites. Carbamidomethylation of cysteine residues was chosen as a fixed modification whereas the N-terminal protein acetylation and oxidation of methionine residues were chosen as variable modifications. The peptide spectrum match (PSM) was set at 1%, as was the false discovery rate (FDR) for proteins, using a target-decoy approach with reversed protein sequences. Perseus (version 1.5.5.3) was used for downstream analysis. The raw data can be found in Proteomics Identification Database (PRIDE) under Project PXD051950.

### Accession numbers

ABCB19 (AT3G28860), ARK1 (AT3G54870), ARK2 (AT1G01950), ARK3 (AT1G12430), BRX (AT1G31880), BRXL2 (AT3G14000), BRXL3 (AT1G54180), BRXL4 (AT5G20540), D6PK (AT5G55910), D6PKL1 (AT4G26610), D6PKL2 (AT5G47750), D6PKL3 (AT3G27580), KIPK (AT3G52890), KIPKL1 (AT2G36350), KIPKL2 (AT5G03640), NEK6 (AT3G44200), PERK8 (AT5G38560), PERK9 (AT1G68690), PERK10 (AT1G26150), PID (AT2G34650), PIN3 (AT1G70940), PIN4 (AT2G01420), PIN7 (AT1G23080), SCR (AT3G54220), ZWI/KCBP (AT5G65930).

## Supporting information

Supplemental Data Set S1

## SUPPLEMENTARY DATA FILES

**Supplementary Table S1:** Primers used in this study.

**Supplementary Data Set S1:** Results from co-immunoprecipitation and related analyses.

## ACKNOWLEDGEMENTS

We thank Christian S. Hardtke (Université de Lausanne, Switzerland), Cyril Zipfel (Universität Zürich, Switzerland), Ronald Pierik (Utrecht University, Netherlands), Shogo Takatani (Nagoya University, Japan) and Jiri Friml (ISTA Klosterneuburg, Austria) for providing seed material. This work was supported by a postdoctoral fellowship from the Alexander-von-Humboldt foundation (NLD 1216207 HFST-P) to YX and grants from the Deutsche Forschungsgemeinschaft to CS (SCHW751/12-2, SCHW751/14-1, SCHW751/15-1, SCHW751/16-1) and UZH (HA3468/6-3). SB and BK received support from the Elitenetzwerk Bayern (F-6-M5613.6.K-NW-2021-411/1/1). JS and KL acknowledge support from the Knut and Alice Wallenberg Foundation (KAW) and the Swedish Research Council (VR), and the Swedish Metabolomics Centre for access to instrumentation.

## AUTHOR CONTRIBUTIONS

YX and CS designed the study, performed all experiments except listed below, performed data analyses and wrote the paper; MZ characterized BRXL2 phosphorylation; AELB supervized biochemical and cell biological analyses; BW performed initial *kipk* and *kipkl* mutant genotyping and phenotyping; JS and KL contributed IAA measurements; DJ, LS and UZH designed and performed oocyte auxin transport experiments; SB and BK performed mass spectrometry and data analysis; JL co-supervized experiments conducted at Guangzhou University. All authors edited and approved the manuscript.

---

**Supplementary Figure S1.**
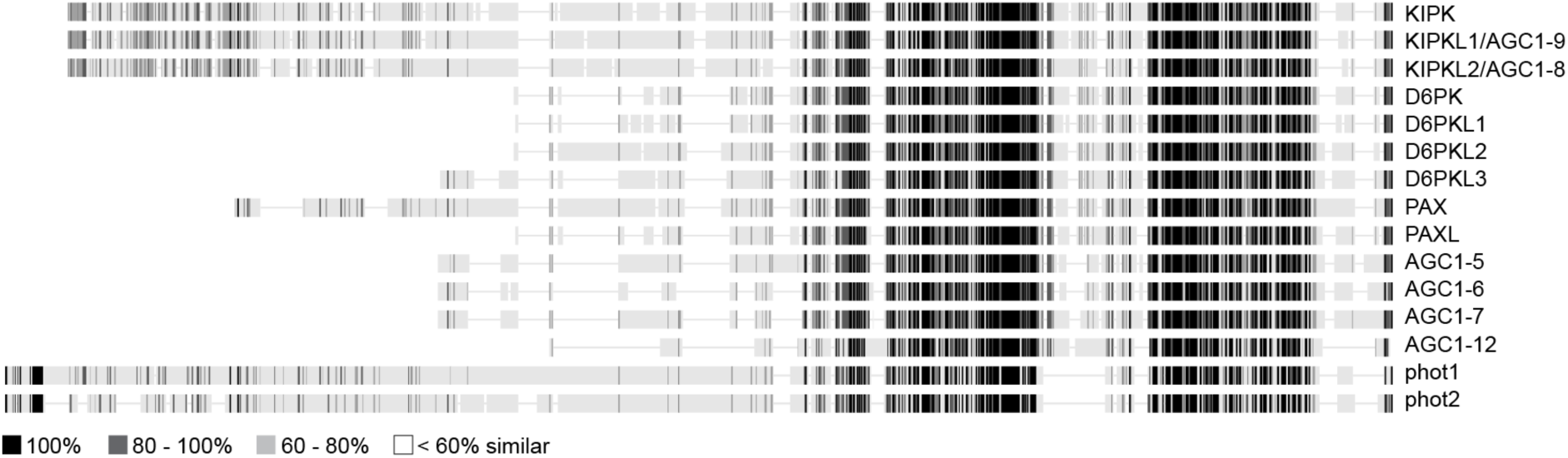
Schematic representation of KIPK and KIPKL proteins in the context of *Arabidopsis thaliana* AGC1 and phototropin AGC4 kinases. Schematic representations of KIPK, KIPKL1/AGC1-9 and KIPKL2/AGC1-8, as well as the remaining 10 AGC1 and the two AGC4 blue light receptor serine/threonine kinases phot1 and phot2 from *Arabidopsis thaliana*. AGC1 and AGC4 kinases are two of four subfamilies of the AGCVIII kinase family.

**Supplementary Figure S2.**
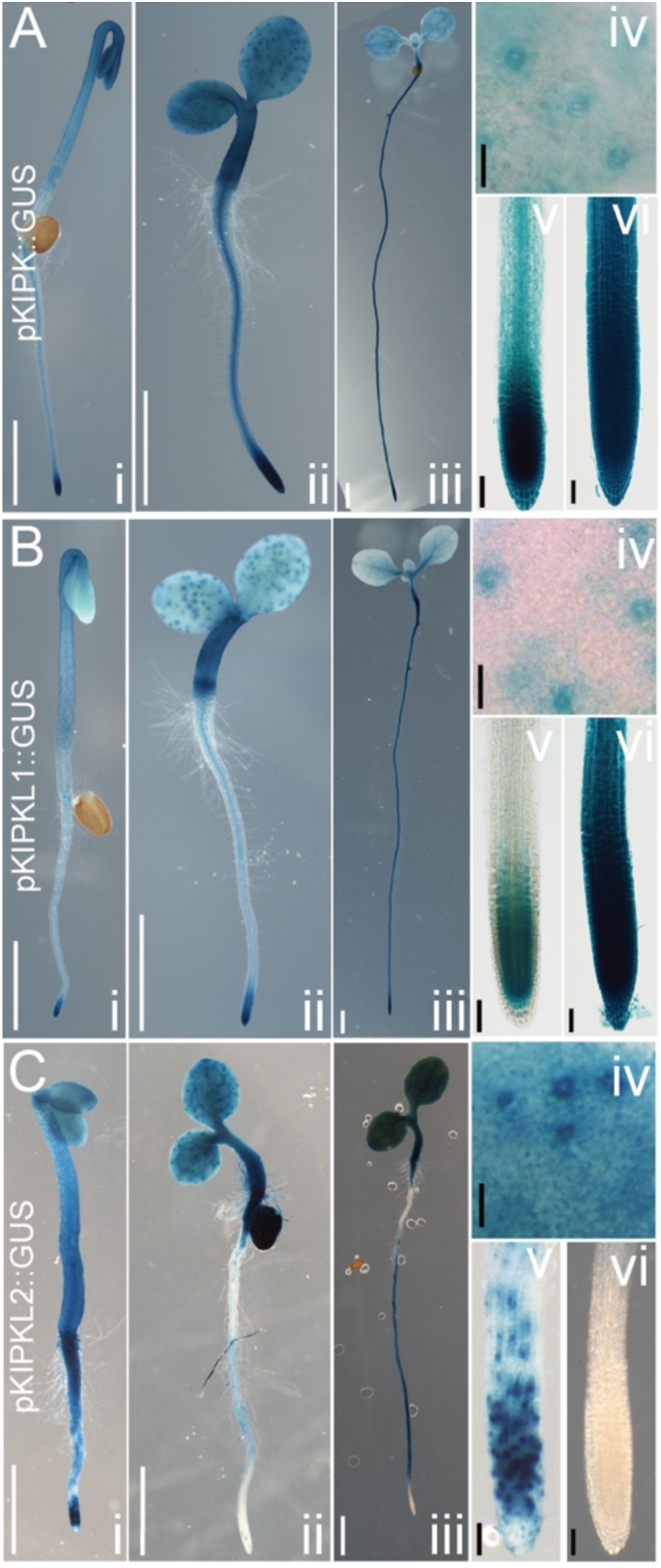
Expression analyses using promoter::GUS fusions reveal a broad expression pattern of the *KIPK* and *KIPKL* genes. **(A)** – **(C)** Photographs of 2.5 days-old dark-grown (i), three- and five-days-old light-grown (ii, iii) seedlings, leaf surfaces with dotted patterns represent stomata staining (iv), as well as root tips of i (v) and iii (vi) from pKIPK::GUS (A), pKIPKL1::GUS (B), and pKIPKL2::GUS (C). Scale bars: 5 mm (i), 3 mm (ii), 3 mm (iii), 40 μm (iv), 50 μm (v), 50 μm (vi).

**Supplementary Figure S3.**
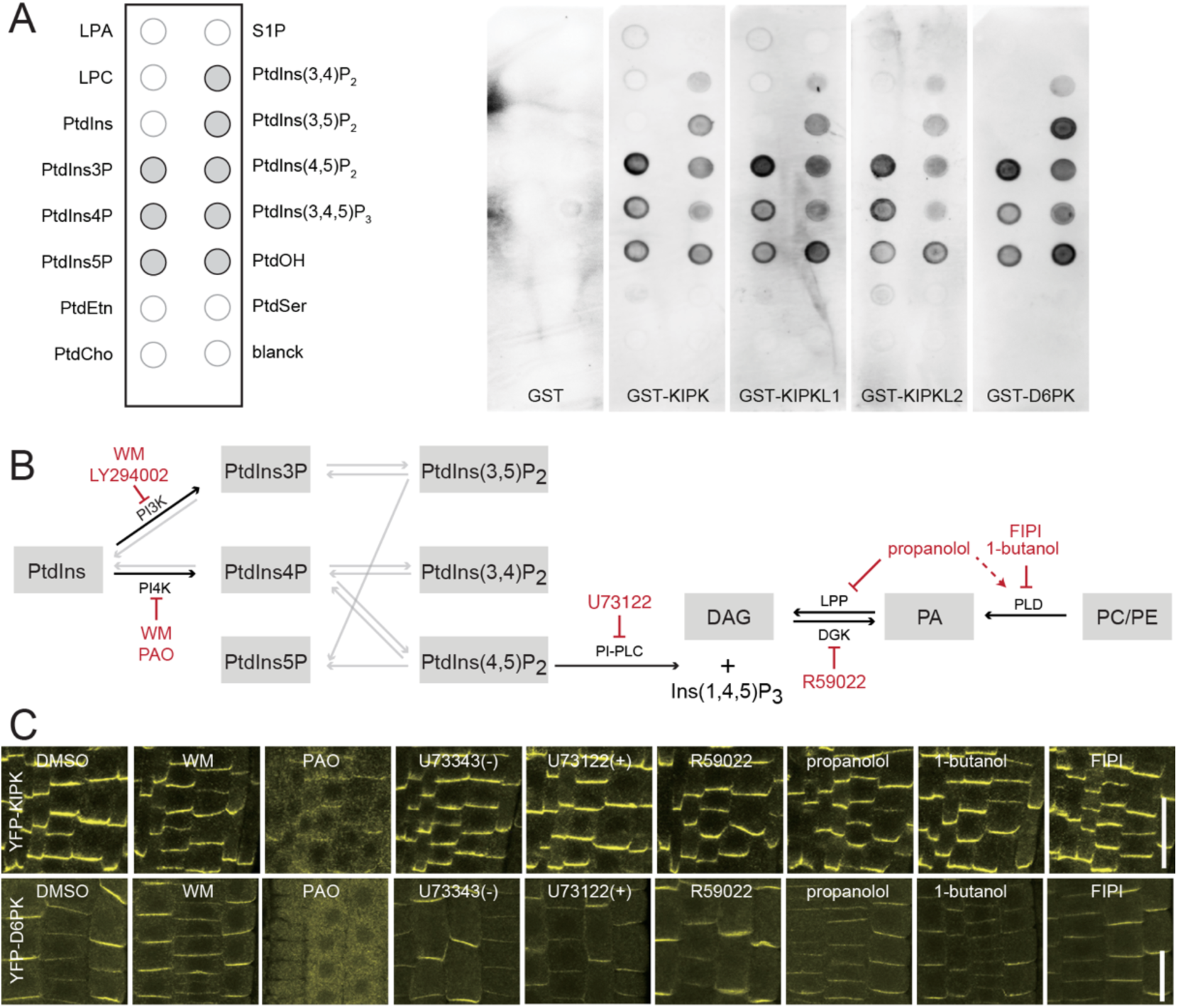
KIPK, KIPKL1 and KIPKL2 bind anionic phospholipids. **(A)** Results of lipid overlay assays with purified GST and GST-tagged kinases as specified. Grey dots in the left panel indicate identities of phospholipids bound by all kinases. **(B)** Schematic overview of the biosynthetic pathways for phosphatidylinositol (PtdIns) and phosphatidic acid (PtdOH) biosynthesis and their chemical inhibitors (red). Black arrows, biosynthetic steps analysed in this study (Meijer and Munnik, 2003; Heilmann, 2009; Potocky et al., 2014; Barbosa et al., 2016; Simon et al., 2016). Abbreviations: DAG, diacylglycerol; DGK, DAG KINASE; LPA, lysophosphatidic acid; LPC, lysophosphatidylcholine; LPP, LIPID PHOSPHATE PHOSPHATASE; PAO, phenylarsine oxide; PI-PLC, PI-SPECIFIC PHOSPHOLIPASE C; PLD, PHOSPHO-LIPASE D; WM, Wortmannin; PtdCho, phosphatidylcholine; PtdEtn, phosphatidylethanolamine; PtdIns, phosphatidylinositol and its mono-/bis-/tris-phosphates; PtdSer, phosphatidylserine; S1P, sphingosine-1-phosphate. **(C)** Representative confocal images of epidermal cells expressing YFP-KIPK or YFP-D6PK after mock treatment (30 min) and treatments (30 min) with the specified inhibitors: 0.1% DMSO, 33 µM WM (Wortmannin), 30 µM PAO (phenylarsenic oxide), 5 µM U73343 (−) inactive and U73122 (+) active analogues, 50 µM R59022, 50 µM propanolol, 0.8% 1-butanol, 1 µM FIPI (5-fluoro-2-indolyl des-chlorohalopemide). The images of the DMSO control and the PAO treatment are identical to the ones shown in Figure 1C. Scale bars = 20 µm.

**Supplementary Figure S4.**
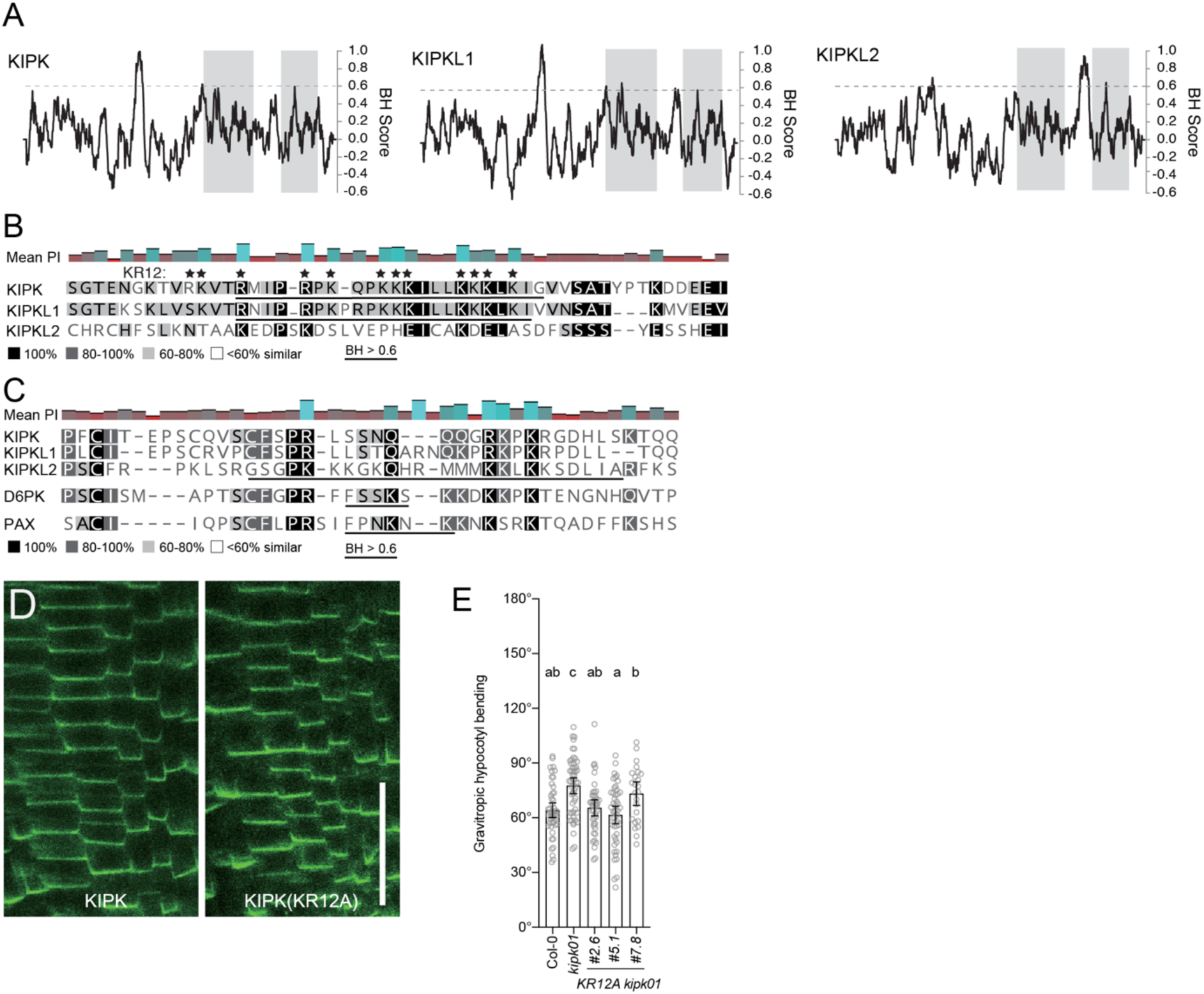
A polybasic region in the N-terminus of KIPK and KIPKL1 is dispensable for plasma membrane interactions. **(A)** Basic hydrophobicity (BH) profiles of KIPK, KIPKL1 and KIPKL2. A BH score greater than 0.6 had previously been shown to be a good predictor for interactions with phospholipids (Bailey and Prehoda, 2015). **(B)** and **(C)** Protein sequence alignments of the basic hydrophobic regions (BH > 0.6) from the N-terminal regions of the specified proteins (B) or from their middle domains (C). Asterisks mark the 12 K and R residues that were mutagenized to A to obtain KIPK(KR12A). **(D)** Representative confocal images of root epidermis cells expressing eGFP-KIPK or eGFP-KIPK(KR12A) in the *kipk01* mutant. Scale bar = 25 µm. **(E)** Graph displaying the average and 95% confidence interval, as well as the individual data points from a negative hypocotyl gravitropism experiment. n > 23 seedlings. Results from a One-way ANOVA analysis are displayed on the top of each bar.

**Supplementary Figure S5.**
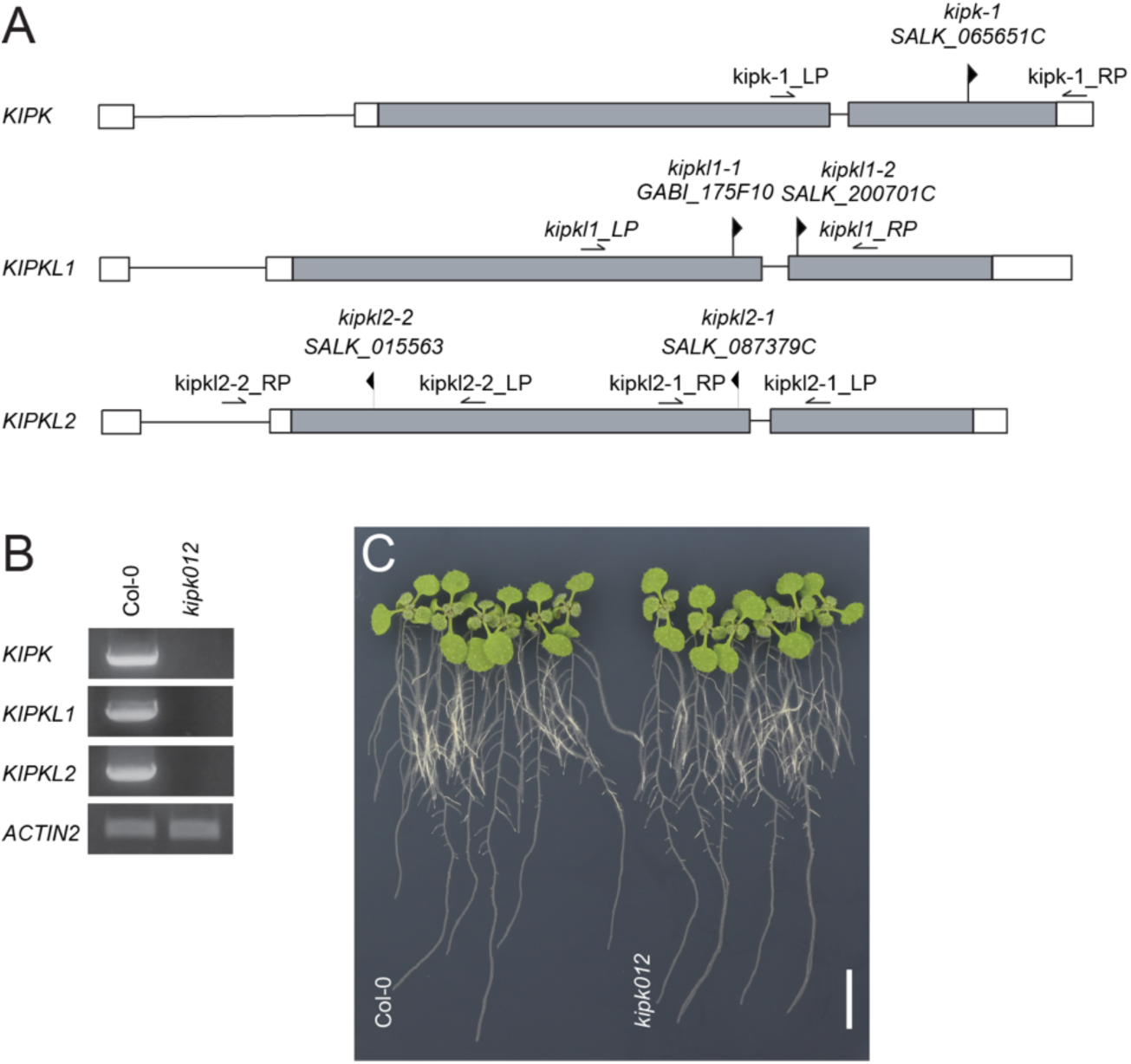
Mutants of *KIPK* and *KIPKLs* do not display apparent growth defects. **(A)** Schematic representation of *Arabidopsis thaliana KIPK*, *KIPKL1* and *KIPKL2* genes with positions of T-DNA insertions in their mutant alleles. The arrowheads indicate the position and the direction of the T-DNA insertion as deposited at the SIGNAL web resource (signal.salk.edu). Left (LP) and right (RP) border primers for genotyping and RT-sqPCR are shown by arrows. **(B)** Results from reverse transcription semi-quantitative PCR (RT-sqPCR) analyses using the primers specified in (A) of the *kipk012* mutant with the alleles *kipk-1, kipkl1-1*, and *kipkl2-1*. *ACTIN2* serves as a control gene transcript. **(C)** Representative photograph of 14-days-old light-grown wild type and *kipk012* plants. Scale bar = 1 cm.

**Supplementary Figure S6.**
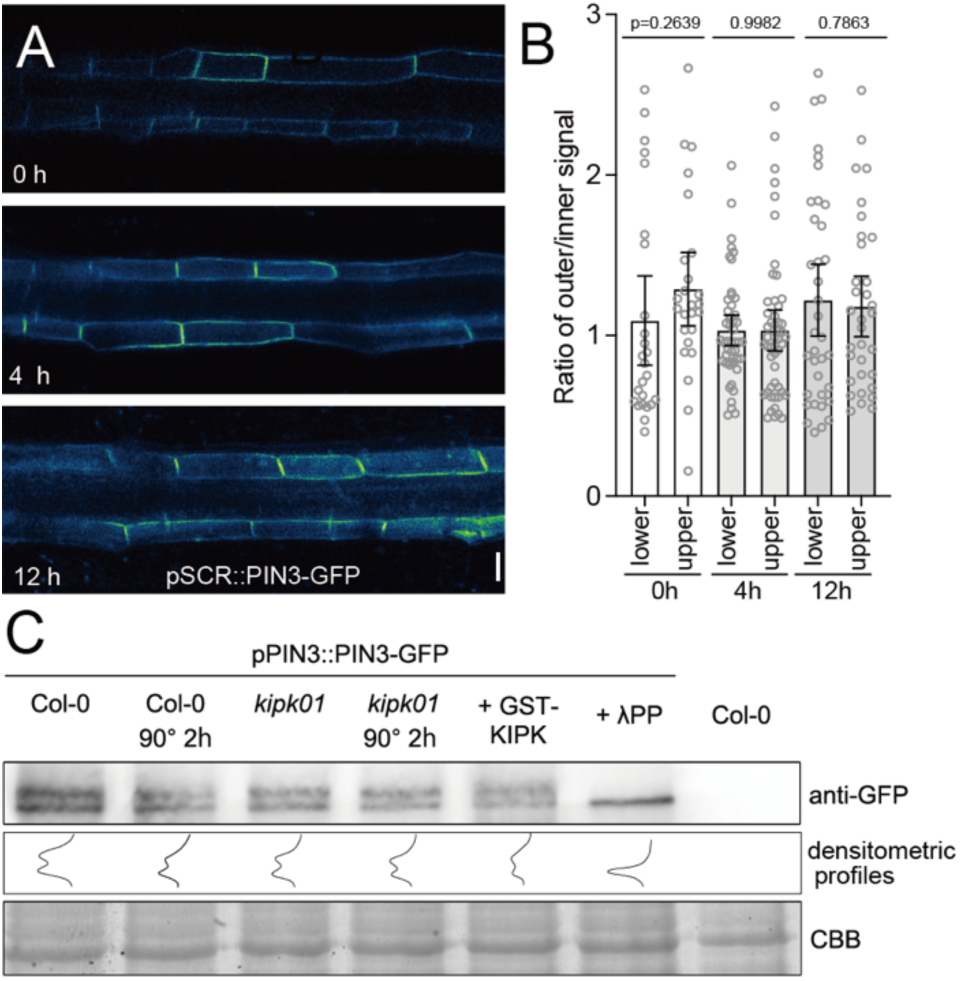
PIN3-GFP lateral distribution is stable during gravitropism response. **(A)** Representative confocal microscopy images of hypocotyl sections of three-days-old dark-grown seedlings expressing pSCR::PIN3-GFP at time points 0, 4 and 12 hrs after gravistimulation. Scale bar = 20 µm. n > 10 seedlings. **(B)** Graph displaying the average and 95% confidence interval, as well as the individual data points of ratios between the outer and inner PIN3-GFP signal during gravitropic hypocotyl bending at time points 0, 4 and 12 hrs after gravistimulation. **(C)** Representative western blot with anti-GFP antibody for the detection of PIN3-GFP before and 2 hrs after gravistimulation in the wild type (Col-0), *kipk01* and after addition of purified recombinant GST-KIPK or l phosphatase (lPP) from four-days-old dark-grown seedlings expressing pPIN3::PIN3-GFP or the non-transgenic wild-type. The upper band corresponds to a phosphorylated form of PIN3-GFP, as revealed by the absence of this band after phosphatase treatment. Densitometric profiles (middle panel) do not suggest major changes in the abundance of PIN3-GFP between the wild type (Col-0)and *kipk01*, or in the phosphorylated PIN3-GFP form between the genotypes and treatments or following gravistimulation. CBB, Coomassie Brilliant Blue-stained gel section, loading control.

**Supplementary Figure S7.**
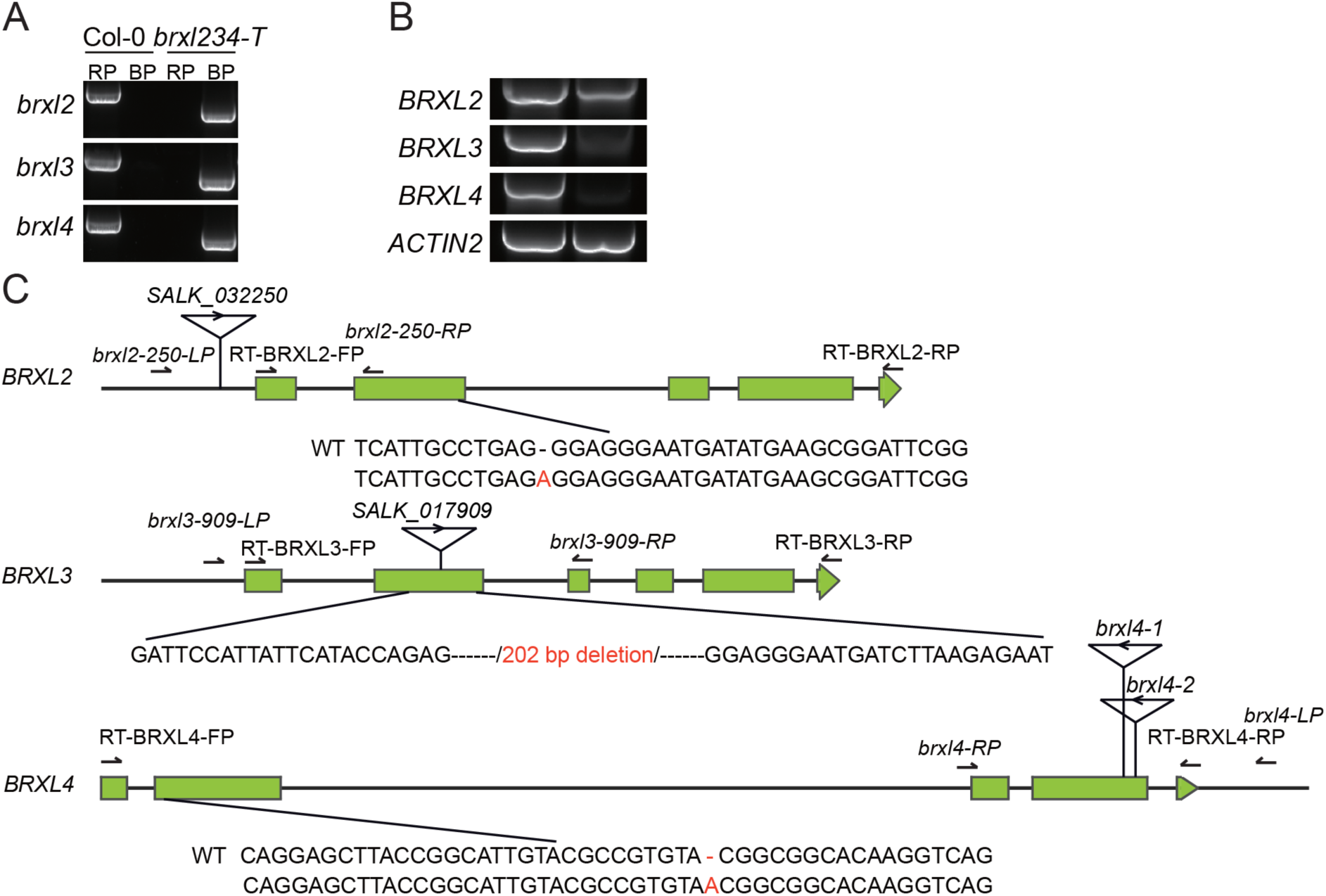
Schematic representation and molecular description of mutant alleles of *BRXL2*, *BRXL3* and *BRXL4*. **(A)** Result of a genotyping analysis of a previously established *brxl2 brxl3 brxl4* (*brxl234-T*) mutant (Briggs et al., 2006). **(B)** Result from a semi-quantitative RT-PCR for *BRXL2*, *BRXL3*, *BRXL4* and the *ACTIN2* control gene of *brxl234-T* reveals substantial expression of *BRXL2*. **(C)** Schematic representation of the exon (boxes) and intron (interrupted lines) structure of *BRXL2*, *BRXL3* and *BRXL4*. Wild type (WT) and mutant allele sequences are shown with CRISPR/Cas9-induced mutations highlighted in red. The positions of T-DNA insertion mutations present in previously described *brxl234-T* with the alleles *brxl4-1* and *brxl4-2*, carrying insertions at the 3’-end of the open reading frame, are also shown (Che et al., 2023).

**Supplementary Figure S8.**
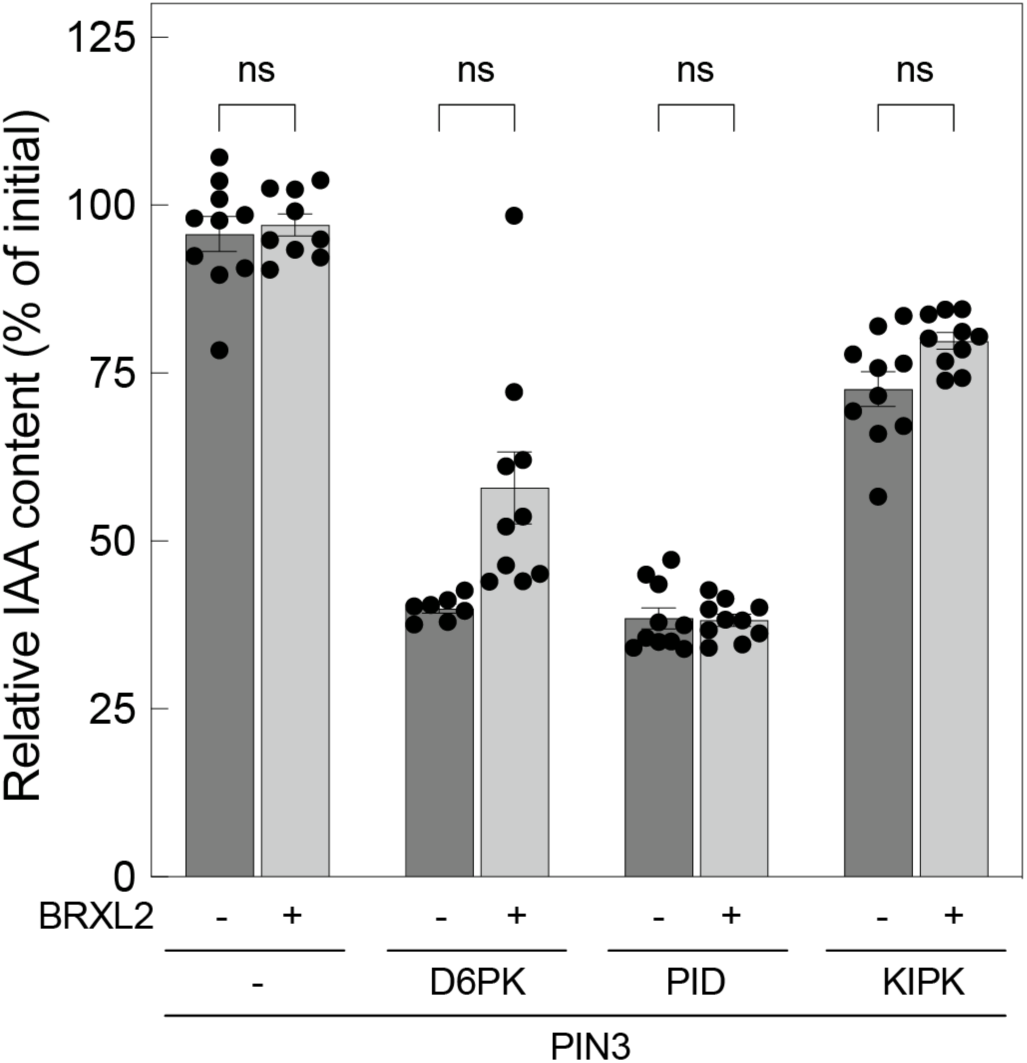
BRXL2 co-expression has no effect on KIPK-mediated PIN3 activation. Results from auxin transport experiments with PIN3, protein kinases and BRXL2 addition as specified. Shown is the IAA content after injection (dark grey bars) set to 1 in comparison to the IAA content after 15 min of efflux (light grey bars) expressed relative to the starting content. Data points represent (10 ≧ n ≧ 9) individual oocytes, bars represent mean and standard error. Groups were compared by a Student’s t-test: ns, not significant.

**Supplementary Figure S9.**
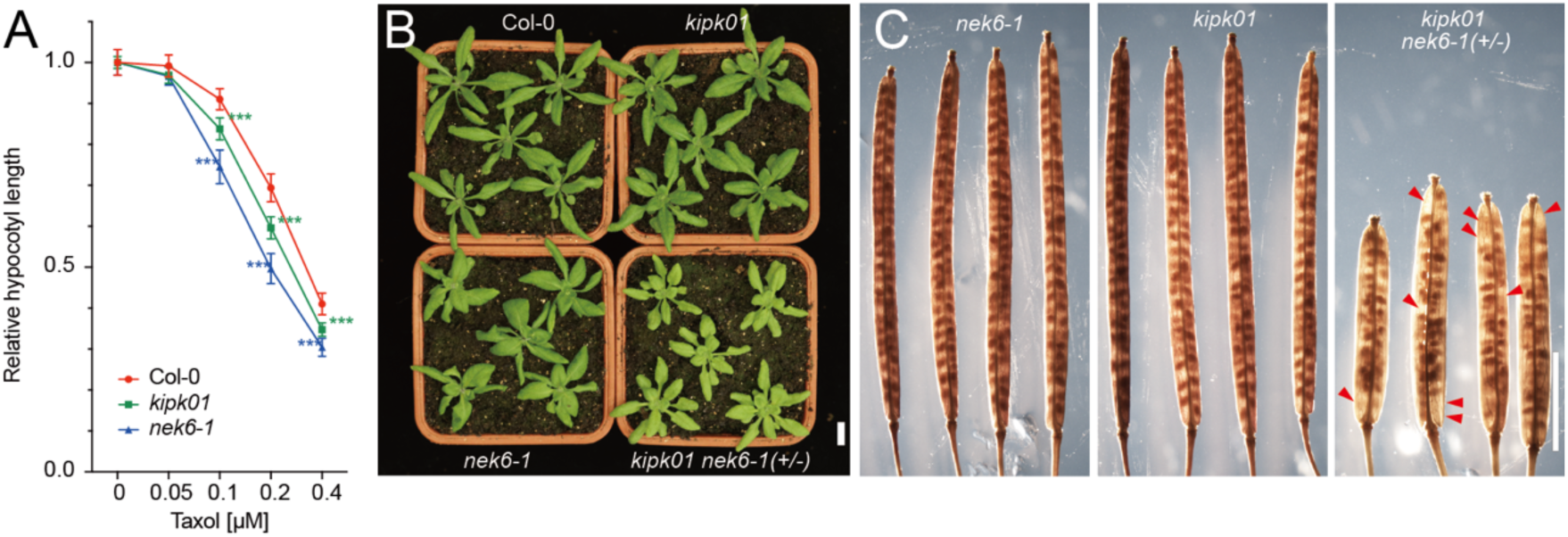
Genetic interaction between *KIPK*, *KIPKL1* and *NEK6*. **(A) (F)** Graph displaying average and 95% confidence interval of relative hypocotyl length measured from four-days-old dark-grown seedlings (n ≥ 55) grown on increasing concentrations of the microtubule-restabling drug taxol. Student’s t-test: *** p < 0.001. **(B)** and **(C)** Representative photographs of 28-days-old wiild type (Col-0), *kipk01*, *nek6-1* and *kipk01 nek6-1* (+/−) plants (B) and mature siliques (C). In (C), red arrowheads mark aborted seeds, a phenotype only apparent in the progeny of *kipk01 nek6-1* (+/−) plants. Scale bars = 1 cm (B) and 4 mm (C).

**Supplementary Figure S10.**
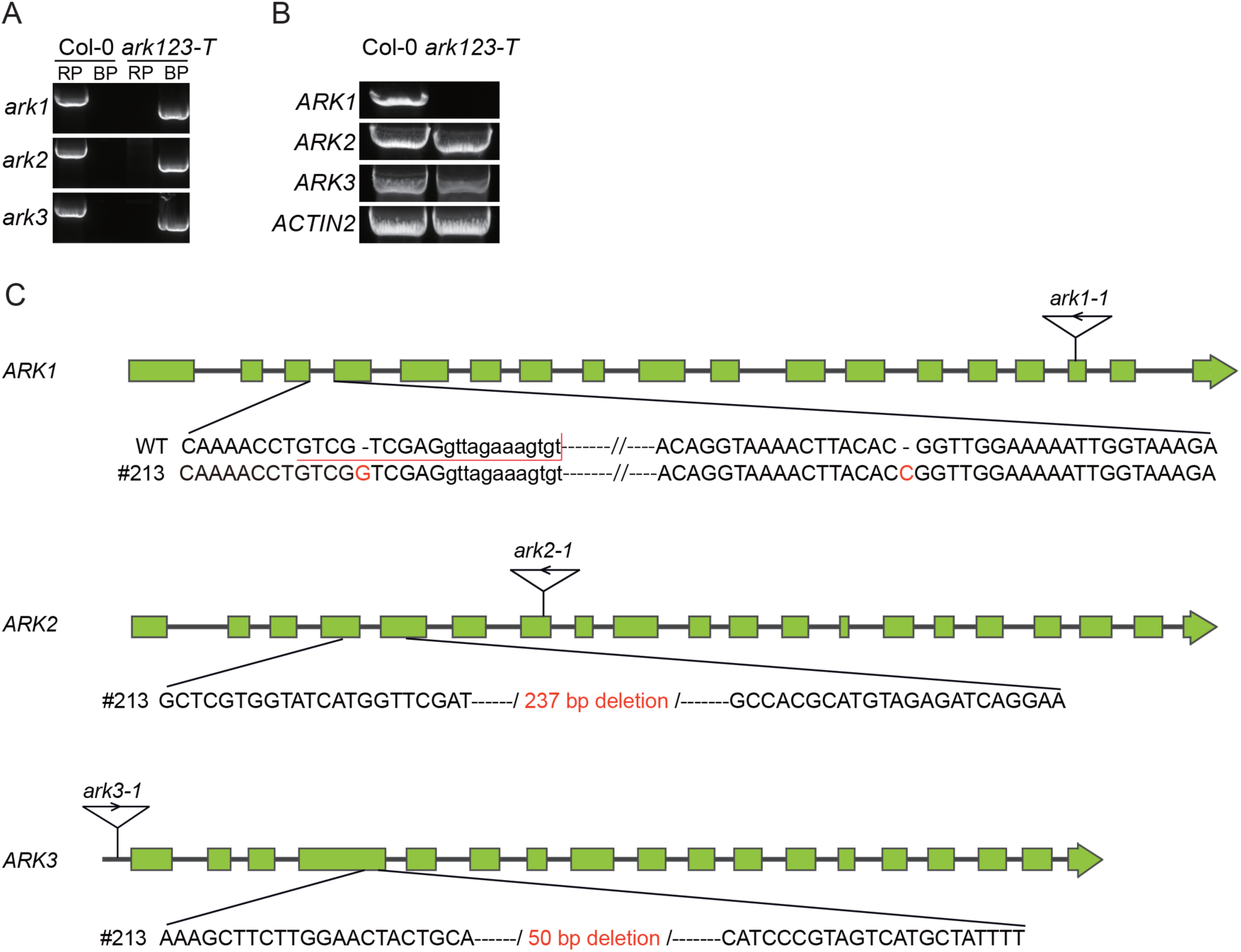
Schematic representation and molecular description of mutant alleles of *ARK1*, *ARK2* and *ARK3*. **(A)** Result of a genotyping analysis of a previously established *ark1-1 ark2-1* and *ark3-1* (*ark123-T*) T-DNA triple mutant (Sakai et al., 2008). **(B)** Results from semi-quantitative RT-qPCR for *ARK1*, *ARK2*, *ARK3* and the *ACTIN2* control gene of *ark123-T* reveals substantial expression of *ARK2 and ARK3* in the mutant. DNA sequencing revealed that ARK2 in *ark2-1* is a truncated version lacking exon 7 carrying the T-DNA insertion. (**C)** Schematic representation of the exon (boxes) and intron (interrupted lines) structure of *ARK1*, *ARK2* and *ARK3*. Wild type (WT) and mutant allele sequences are shown with CRISPR/Cas9-induced mutations highlighted in red. T-DNA insertions present in the previously *ark1-1 ark2-1* and *ark3-1* (*ark123-T*) triple mutant are also indicated (Sakai et al., 2008).

**Supplementary Figure S11.**
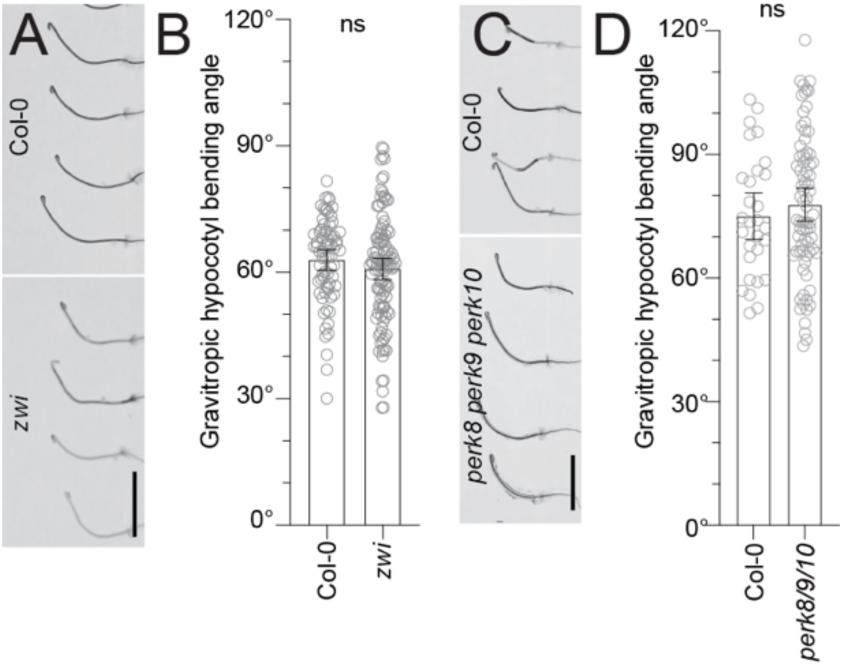
*ZWI* and *PERK8 PERK9 PERK10* are not required for gravitropic hypocotyl bending. **(A)** and **(C)** Representative photographs of three-days-old dark-grown seedlings of the specified genotypes 24 hours after reorientation by 90°. Scale bars = 1 cm. **(B)** and **(D)** Graphs displaying the average and 95% confidence interval, as well as the individual data points (n ≥ 55) from a negative hypocotyl gravitropism experiment as shown in (A) and (B). Student’s t-test: ns, not significant.

